# A new massively-parallel transposon mutagenesis approach comparing multiple datasets identifies novel mechanisms of action and resistance to triclosan

**DOI:** 10.1101/596833

**Authors:** Muhammad Yasir, A. Keith Turner, Sarah Bastkowski, Andrew J. Page, Andrea Telatin, Minh-Duy Phan, Leigh G. Monahan, Aaron E. Darling, Mark A. Webber, Ian G. Charles

**Affiliations:** Quadram Institute Bioscience, Norwich Research Park, Norwich, NR4 7UQ, UK; School of Chemistry and Molecular Biosciences, The University of Queensland, St Lucia 4072, Queensland, Australia; The ithree institute, University of Technology Sydney, PO Box 123, Broadway NSW 2007, Australia; University of East Anglia, Norwich Research Park, Norwich, NR4 7TJ

**Keywords:** TraDIS, fatty acid biosynthesis, antibiotic resistance, efflux, bioinformatics

## Abstract

The mechanisms by which antimicrobials exert inhibitory effects against bacterial cells and by which bacteria display resistance vary under different conditions. Our understanding of the full complement of genes which can influence sensitivity to many antimicrobials is limited and often informed by experiments completed in a small set of exposure conditions. Capturing a broader suite of genes which contribute to survival under antimicrobial stress will improve our understanding of how antimicrobials work and how resistance can evolve. Here, we apply a new version of ‘TraDIS’ (Transposon Directed Insert Sequencing); a massively parallel transposon mutagenesis approach to identify different responses to the common biocide triclosan across a 125-fold range of concentrations. We have developed a new bioinformatic tool ‘AlbaTraDIS’ allowing both predictions of the impacts of individual transposon inserts on gene function to be made and comparisons across multiple TraDIS data sets. This new TraDIS approach allows essential genes as well as non-essential genes to be assayed for their contribution to bacterial survival and growth by modulating their expression. Our results demonstrate that different sets of genes are involved in survival following exposure to triclosan under a wide range of concentrations spanning bacteriostatic to bactericidal. The identified genes include those previously reported to have a role in triclosan resistance as well as a new set of genes not previously implicated in triclosan sensitivity. Amongst these novel genes are those involved in barrier function, small molecule uptake and integrity of transcription and translation. These data provide new insights into potential routes of triclosan entry and bactericidal mechanisms of action. Our data also helps to put recent work which has demonstrated the ubiquitous nature of triclosan in people and the built environment into context in terms of how different triclosan exposures may influence evolution of bacteria. We anticipate the approach we show here that allows comparisons across multiple experimental conditions of TraDIS data will be a starting point for future work examining how different drug conditions impact bacterial survival mechanisms.

## Introduction

Triclosan is a broad-spectrum antimicrobial which was first developed in the 1960s. Since then it has been used extensively in a wide range of products for clinical, veterinary and domestic use^1^, such as hand soaps, mouthwashes, toothpaste, shampoos and cosmetics as well as being incorporated into plastics. Triclosan acts by inhibiting type II fatty acid biosynthesis and is the archetypal inhibitor of this essential pathway in bacteria. Over the years, other compounds that inhibit enzymes in this pathway have progressed to different stages of clinical development as novel antibiotics, but triclosan is the only one in widespread domestic use. Triclosan specifically inhibits the enzyme FabI; an enoyl-acyl carrier protein (ACP) reductase conserved across bacteria, hence triclosan has a correspondingly broad spectrum of activity against most bacteria^2^.

The use of triclosan has proliferated to such an extent that it is readily identified in most aquatic environments and is detectable in urine and blood plasma of most people in the developed world where use has been extensive in personal care products, particularly toothpastes^3–5^. A US study found approximately 10% of adults had levels of triclosan in their urine above the minimum inhibitory concentration required to inhibit most bacteria^6^.

There has been pressure to ban triclosan use due to concerns about its potential impact as a hormone-disrupting agent and evidence that exposure to triclosan can select for mutants cross-resistant to antibiotics^7–9^. Evidence from mouse studies has suggested that triclosan exposure can alter composition of the gut microbiota and may induce inflammation and promote colonic cancers^10^. The impact of triclosan in the built environment, where it is now a common contaminant in developed countries^11^, has recently been investigated. Correlations between triclosan concentrations and organisms present in dust strongly suggest the selection of drug-resistant bacterial communities in the presence of triclosan^11^.

Whilst the primary target of triclosan has been established as FabI, a full understanding of triclosan mechanisms of action has remained elusive. It has been demonstrated that triclosan exerts a bacteriostatic effect at low concentrations but at higher concentrations is bactericidal^12^. Triclosan resistance also appears multifactorial; highly resistant mutants typically have substitutions within FabI which reduces binding efficiency of triclosan, but multidrug efflux pumps, barrier function, changes to *fabI* expression, core metabolic pathways and sigma factor-controlled stress responses have all been implicated as being involved in triclosan resistance^13–17^. A specific link between resistance to fluoroquinolone antibiotics and triclosan, mediated by altered stress responses present in *gyrA* mutants, has also recently been described^8^.

Given renewed interest in the impact of triclosan on population health and the microbiome we felt it was timely to systematically investigate the mechanisms of action and resistance of triclosan across a range spanning bacteriostatic and bactericidal concentrations. This will help understand the potential impacts and risks of exposure to triclosan under different conditions as well as informing fatty acid inhibitor development programmes and providing further insight into the impacts of triclosan on the microbiome.

Previous systems biology approaches to investigate bacterial responses to triclosan have focused on nucleotide sequencing the genomes of mutants, transcriptomics and proteomics analysis which have tended to use a small number of concentrations^13, 16^. These studies have been informative although each is limited to providing a snapshot of genes potentially involved in triclosan susceptibility under one condition.

Here we applied a modified version of ‘TraDIS’ (<u>Tra</u>nsposon <u>D</u>irected Insertion-site <u>S</u>equencing, ^18, 19^) to allow conditional fitness of all genes in the genome to be assayed simultaneously, including essential genes which cannot effectively be assayed in traditional TraDIS experiments. Our new methodology uses a transposon with an outward-directed inducible promoter allowing the impact of gene over-expression and gene silencing of every gene in a candidate organism to be assayed (in addition to insertional inactivation) and we refer to this approach as TraDIS+. Using TraDIS+ we analysed the response of *Escherichia coli* to triclosan at eight concentrations spanning a 125-fold range. We also employed a refined analysis pipeline we have developed (AlbaTraDIS) able to identify and compare changes in insert site abundance from multiple exposure conditions as well as predicting impacts of insertions on gene expression automatically.

TraDIS+ builds on previous studies that have systematically attempted to artificially over-express known antibiotic drug targets to assess their impact on susceptibility. The method effectively integrates a highly regulated promoter (at high density) throughout a target genome allowing changes in survival and growth to be assessed for all genes without prior knowledge of their function. The ability to regulate expression of all the genes of a target organism in one massively-parallel experiment in a controlled manner promises to be a hugely powerful strategy that has the potential to uncover many new genotype-phenotype relationships.

Using this approach, we clearly show distinct concentration-dependent responses to triclosan and identify novel genes involved in triclosan susceptibility including identification of potential routes of entry into the cell. We were also able to systematically document different impacts between exposure to bacteriostatic and bactericidal concentrations of the drug for the first time.

## Results

### Creation of a high-density transposon insert library in E. coli strain BW25113

*E. coli* BW25113 was transformed with a bespoke Tn5-based transposon containing an outward facing *tac* promoter. To confirm that a suitable number of unique transposon inserts had been harvested and that the modified TraDIS sequencing strategy worked, DNA was extracted from the transposon mutant library pool and was processed for TraDIS sequencing (**Supplementary figure 1**). In this experiment, 5.1 million TraDIS reads were mapped to the BW25113 reference genome at 813873 unique sites, giving an average of one insertion site every 6 bp (**Supplementary figure 2**). Analysis of the insert sites revealed an expected lack of inserts in genes previously reported as essential^20^. These data confirmed a high-density mutant library covering the genome had been created, that the modified TraDIS sequencing method worked as expected and that non-specific reads were not being generated. We also assessed the reproducibility of experiments by comparing data from two separate cultures seeded from the library by sequencing transposon insert sites in control conditions and in the presence of three concentrations of triclosan (**Supplementary figure 2**). This data showed excellent correlation between the replicates. Finally, 48 individual transposon mutants were randomly selected and whole genome sequenced to check insert sites. Bioinformatic analysis of these mutants confirmed that each mutant contained only one copy of the transposon inserted in the genome at unique positions, thus supporting the assertion that the transposon mutant library pool consists largely of mutants with single transposon insertions.

### TraDIS+ is a powerful tool which identifies multiple genes involved in triclosan sensitivity

A total of 66 independent TraDIS experiments were completed across a 125-fold concentration range of triclosan. Analysis of differential abundance of insert sites in the presence of triclosan identified previously reported mechanisms of triclosan resistance as highly significant targets. FabI is the primary target for triclosan: an essential gene and, as such, it cannot be inactivated by transposon insertion. Our data (**Figure 1**) revealed a highly significant enrichment of transposon inserts upstream of *fabI* in triclosan-treated experiments with an almost complete bias for transposon insertions oriented such that the outward directed promoter will up-regulate transcription of *fabI*. This confirmed that we were able to identify known mechanisms of triclosan resistance and shows that TraDIS+ can readily identify the contribution of an essential gene to survival following exposure to a target drug.

**Figure 1.**
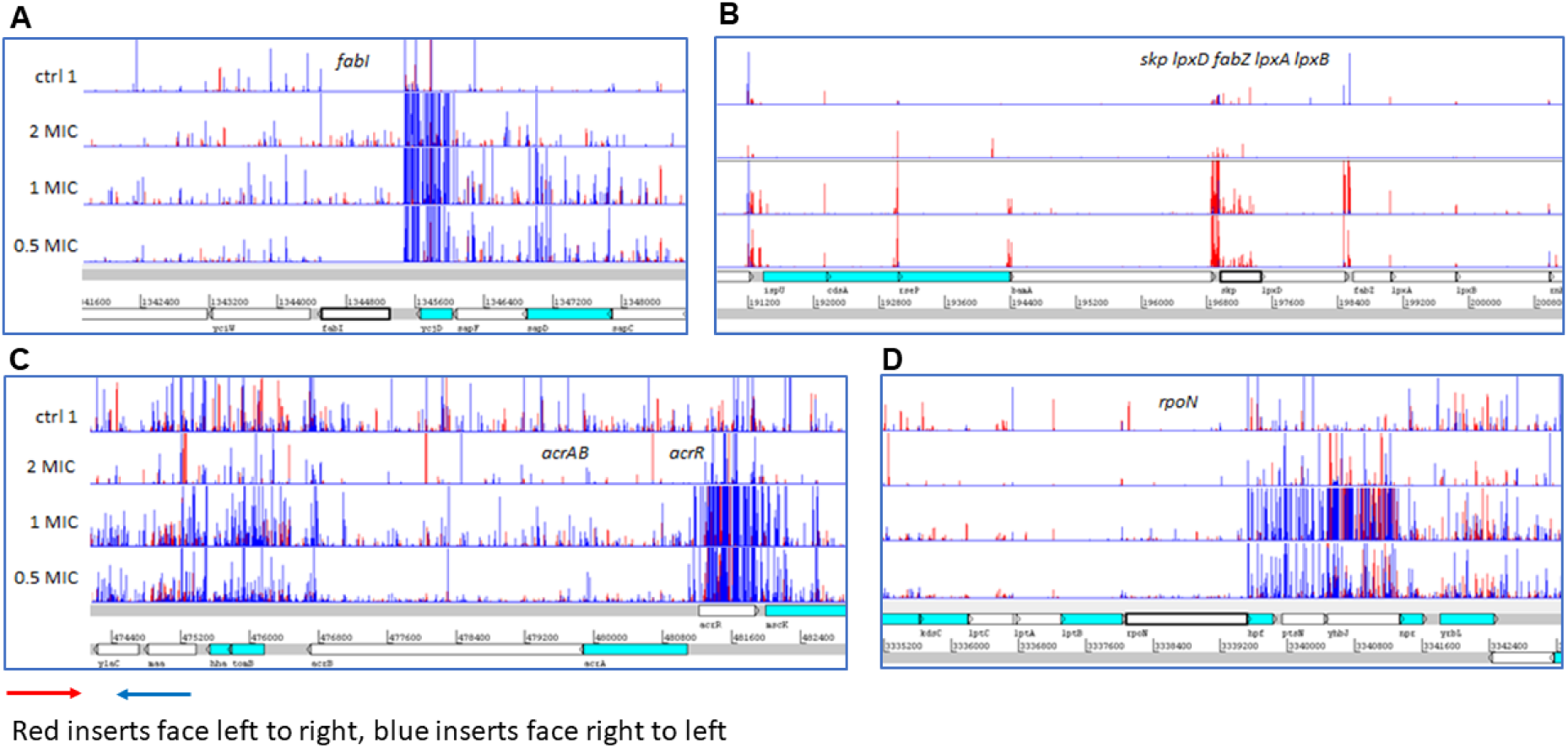
Identification of known targets for triclosan including essential genes. Identification of known targets using the TraDIS+ approach incorporating an inducible outward-facing promoter which identifies the impact of both essential and non-essential genes on survival and growth. A genetic map of the relative gene positions is shown at the bottom of the panel. Above this, each row of vertical red or blue lines (plotted with red behind) indicates the position of mapped reads and the height of the bar represents the relative number of reads mapped. Red indicates transposon insertions wherein the transposon-encoded outward facing promoter is oriented 5’ to 3’ left to right, and blue indicates the opposite right to left orientation. **Panel A** shows inserts predicted to result in up-regulation of *fabI* in the presence of triclosan. **Panel B** shows mutants up-regulating *skp*, *lpxD*, *fabZ*, *lpxA* and *lpxB*. **Panel C** shows mutants inactivating *acrR* and up-regulating *acrAB* being enriched by triclosan. **Panel D** shows mutants positioned antisense to *rpoN* being selected in the presence of triclosan. The top row in each plot shows untreated control cultures, the three rows below are for cultures grown in the presence of 2X, 1X, and 0.5X the MIC of triclosan.

**Figure 2.**
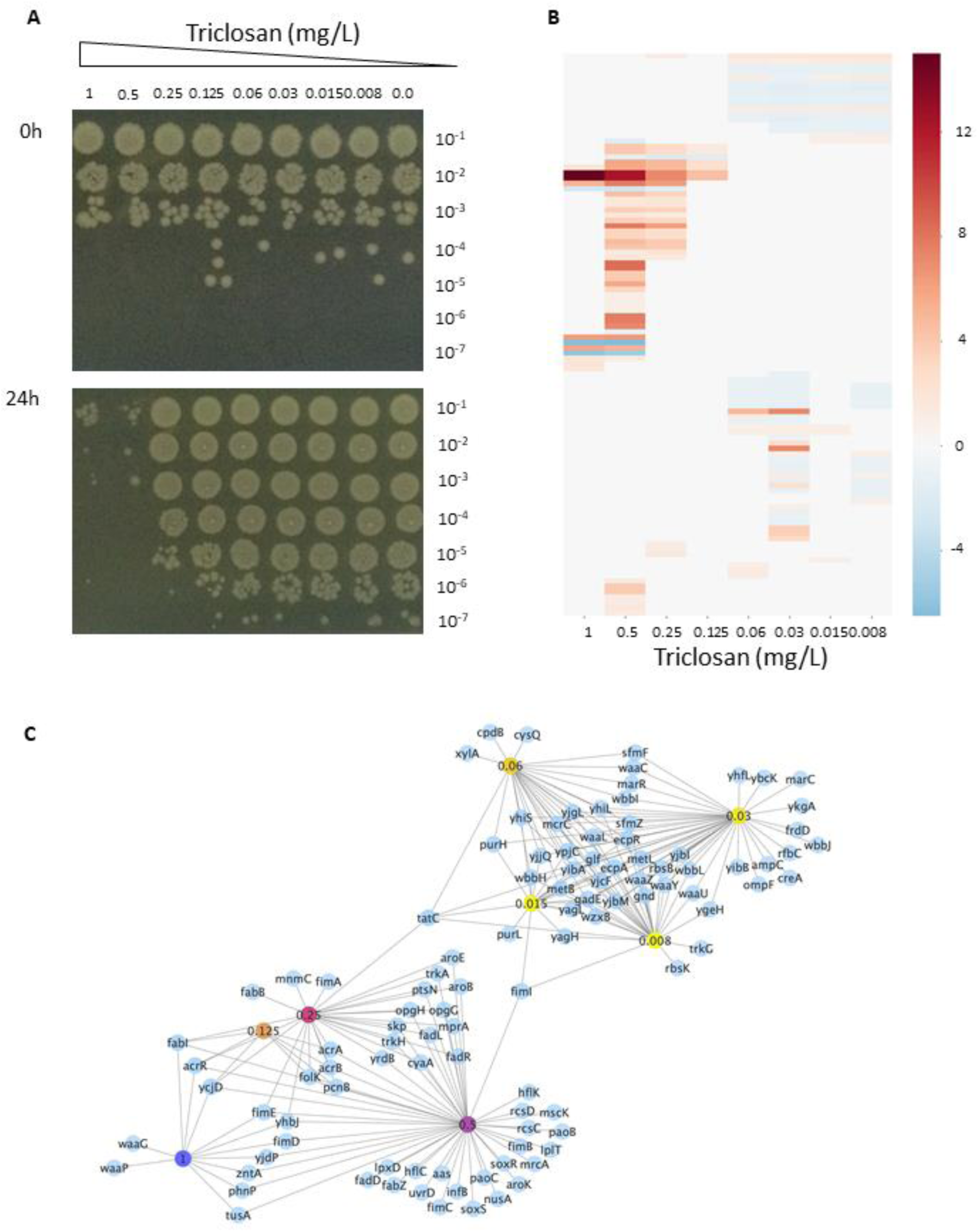
Concentration dependent TraDIS responses and cellular viability. **Panel A** shows the viability of *E. coli* BW 25113 in samples taken from LB broth cultures supplemented with different concentrations of triclosan, immediately after preparation of the broth cultures (top photograph), and after incubation at 37°C for 24 h (bottom photograph). The samples taken from the cultures were serially diluted 10-fold, and 5 µl of all dilutions were spotted onto LB agar and incubated at 37°C overnight to allow surviving bacteria to grow into colonies (shown). The level of dilution (from 10^−1^ to 10^−7^) is shown on the right side of the photographs and the concentrations of triclosan in the cultures are shown along the top (from 1 to 0.008 mg/L; 0.0, no triclosan added). These results show a bactericidal effect above 0.5 mg/L; triclosan at 0.25 mg/L caused a 10-fold reduction in growth over the 24 h period, but at 0.125 mg/L and below there was no obvious growth inhibition. **Panel B** shows a heat map highlighting the differences in sequence reads mapped to genes following growth of the transposon mutant library at the different triclosan concentrations shown along the bottom of the chart. Each row represents a gene and the different genes are ordered from top to bottom. The intensity scale reflecting the size of difference is shown on the right of the map. Some genes show a big difference (darker shades) at several different concentrations of triclosan, while others show more subtle differences (paler shades) at fewer or only one concentration. Read count is a surrogate measure of the number of mutants in the mutant pool, therefore darker shades indicate that mutations in those genes conferred more protection against triclosan. **Panel C** shows a network illustrating the clustering of genes identified as important at different concentrations of triclosan (indicated by coloured nodes). The similarities of responses between sub MIC and supra MIC exposures are clearly visible.

Multidrug efflux pumps and outer membrane porins have also been shown previously to influence triclosan sensitivity including the multidrug efflux system AcrAB. Mutants with inserts within the genes encoding the structural components of this pump were depleted in the presence of triclosan concentrations above the MIC (**Figure 1**) whereas inserts within the local repressor of the system, *acrR* were enriched. Similarly, inserts within outer membrane porin, *ompF* were enriched in the presence of triclosan. Expression of *acrAB and ompF* are controlled by global stress response systems *marRA* and *soxRS*; inserts within *marR* and *soxR* (both of which result in de-repression of the cognate transcriptional activator genes which then promote pump expression) were significantly enriched upon triclosan exposure. Inactivation of sigma factors has also been associated previously with triclosan resistance^14^, in our data there was significant enrichment of inserts immediately downstream of *rpoN* (**Figure 1**) but in an antisense orientation in the presence of triclosan showing that inducible promoters can also identify genes where repression is beneficial under a given stress.

### Exposure to triclosan revealed concentration dependent responses

Whilst the well-established mechanisms of triclosan resistance were identified in our data there were also a large number of genes identified which have not previously been implicated in triclosan resistance. The genes identified were concentration dependent and exposure to different concentrations of triclosan resulted in a range of impacts on cellular viability (**Figure 2**). Similarly, distinct patterns of transposon mutants were revealed across the range of drug exposures. Two major groups of responses were evident from analysis of the data; one block of genes was selected by exposure to concentrations of triclosan at the MIC and below and another block selected by exposure to concentrations above the MIC (**Figure 2, Table 1 and Supplementary Table 1**). Genes involved in fimbrae biosynthesis (*fimABCDEI*) were selected across a wide range of concentrations.

**Table 1.** Selected pathways involved in triclosan sensitivity at different conditions

| Loci/pathway | Function | Predicted impact and change in growth capacity of knock out + triclosan |
| --- | --- | --- |
| **'Sub-MIC' block** |  |  |
| Purine biosynthesis<br>(*purL*, *purH*) | Nucleic acid production | Genes involved in production of nucleotides |
| Lipopolysaccharide biosynthesis<br>(*waaCGLPUYZ*,<br>*wbbHIJL*, *wzxB*) | Barrier function | Genes involved in biosynthesis of lipopolysaccharide protected in the presence of triclosan |
| Methionine biosynthesis<br>(*metBL*) | Amino acid synthesis |  |
| Stress response (*marR*) | Regulation of stress response | Inactivation of *marR*; the repressor of *marA* which co-ordinates an efflux-based stress response |
| **'MIC and above' block** |  |  |
| Stress response (*soxR*) | Regulation of stress response | Inactivation of *soxR*; the repressor of *soxS* which co-ordinates an efflux-based stress response |
| Ribose uptake (*rbsBK*) | Sugar import | Inactivation of the importer selected by triclosan |
| MDR efflux pumps<br>(*acrAB*, *emrA*) | Transport | Multidrug efflux pumps protected in the presence of triclosan |
| Stress response (*rpoN*) | Regulation | Inserts antisense of RpoN enriched in the presence of triclosan |
| Fatty acid biosynthesis (*fabBIZ*) | Membrane barrier | Over-expression of fatty acid biosynthesis selected by triclosan exposure |
| Phospholipid recycling (*mlaBCDEF*) | Membrane barrier | Phospholipid recycling machinery protected in the presence of triclosan |
| Potassium import (*trkAH*) | K<sup>+</sup> importer | Importer disrupted in the presence of triclosan |
| mRNA stability (*pcnB*) | Transcription fidelity | Inactivation of polyA polymerase selected by triclosan |
| Protein stability (*lon*, *infB*, *rsmB*) | Translation fidelity | Disruption of *lon* and *rsmB*, up-regulation of *infB* selected by triclosan |
| Ubiquinones (*ubiBDEFHI*) | Energy generation | Inactivation of ubiquinone biosynthesis selected by triclosan |

The ‘MIC and below’ block was characterised by the protection of genes encoding lipopolysaccharide biosynthesis, purine biosynthesis and methionine biosynthesis. Inactivation of *marR*, the repressor of the global stress response regulator, *marA* was also selected in this block.

The ‘above MIC’ block included mutants which result in de-repression of the global stress regulator, *soxS* which (as with *marA*) controls expression of *acrAB* as well as repression of the major porin, *ompF*. Together these will result in reduced permeability of the cell to triclosan. This block also identified protection of the structural components of the AcrAB multidrug efflux system and over-expression of another efflux system, EmrAB. In addition, inactivation of the ribose uptake system RbsACB and potassium uptake transporter TrkAH were also predicted to be protective of exposure to triclosan. Other loci identified as important for survival at the highest concentrations of triclosan included fatty acid biosynthetic genes (including *fabI*), the phospholipid recycling pathway (*mlaBCDEF*), ubiquitins (*ubiDEFGHL*) and *rpoN*. Additionally, a set of genes all involved in the stability of mRNA, proteins and translation initiation (*pcnB, infB, lon* and *rsmB*) were also identified as contributing to survival at high triclosan concentrations (**Table 1**).

### Triclosan sensitivity is impacted by ribose and potassium import systems

Whilst triclosan has a validated intracellular target in FabI, it is currently unclear how it crosses the cytoplasmic membrane. TraDIS+ data identified inactivation of *rbsB*, the periplasmic ribose binding domain of the RbsABC ribose importer and *trkHA*, components of a potassium importer as being beneficial to survival in the presence of triclosan. Defined *rbsB*, *trkH* or *trkA* mutants from the KEIO mutant collection showed significantly (p <0.01) better growth compared to BW25113 in the presence of 0.125 mg/L of triclosan, although none were able to grow at 0.5 mg/L (**Figure 3**). This data supports the hypothesis these transporters are involved in triclosan sensitivity.

**Figure 3.**
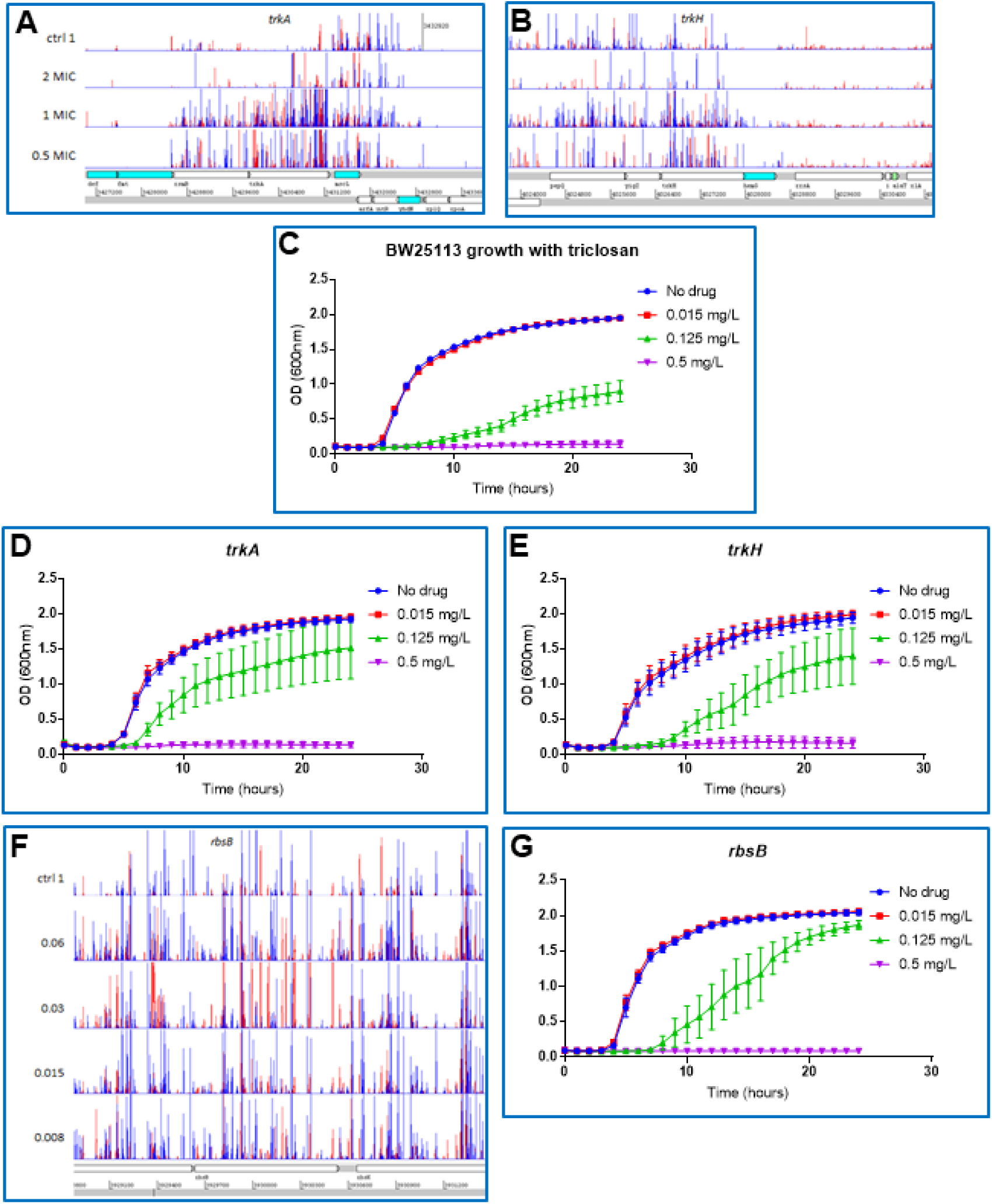
Inactivation of *rbsB* or *trkHA* provides protection against triclosan. **Panels A** and **B** show insert patterns indicating inactivation of *trkAH* is beneficial for survival in the presence of triclosan. **Panels C-E** show growth curves for BW25113 and isogenic mutants grown in LB broth supplemented with different concentrations of triclosan. Panel **F** shows increase in insertions within the first part of *rbsB* in triclosan-exposed libraries. Panel **G** shows growth of a *rbsB* mutant in the presence of triclosan. Growth kinetics data show mean optical density values from multiple replicates (n = 34 for BW25113 and n = 4 for other mutants). Error bars indicate standard error from the mean. The format of the genetic maps is as described in Figure 1.

### Genes maintaining transcript and protein stability are involved in triclosan sensitivity

The mechanism by which triclosan exerts a bactericidal effect at high concentrations is not understood. TraDIS+ identified a set of genes which are involved in mRNA stability, initiation of translation and protein stability as being important to survival in the presence of high concentrations of triclosan. These included *pcnB,* which adds polyA tails to newly synthesised mRNA transcripts that targets their digestion; inactivation of *pcnB* gave a selective advantage during growth in triclosan (**Figure 4**). TraDIS also identified a selective advantage in triclosan for mutants with insertions immediately upstream of both *infB* and *rsmB,* which code for a translation initiation factor and a methyltransferase acting to stabilise binding of initiator tRNAs to the ribosome pre-initiation complex, respectively. Finally, mutants with inactivation of the Lon protease were found to survive better in the presence of triclosan (**Figure 4**). Lon is a general protease (previously implicated in processing of AcrB). These observations suggest that exposure to higher triclosan concentrations impairs mRNA transcript stability and protein translation; thus, mutations that result in increased stability of mRNA and proteins provide a selective advantage.

**Figure 4.**
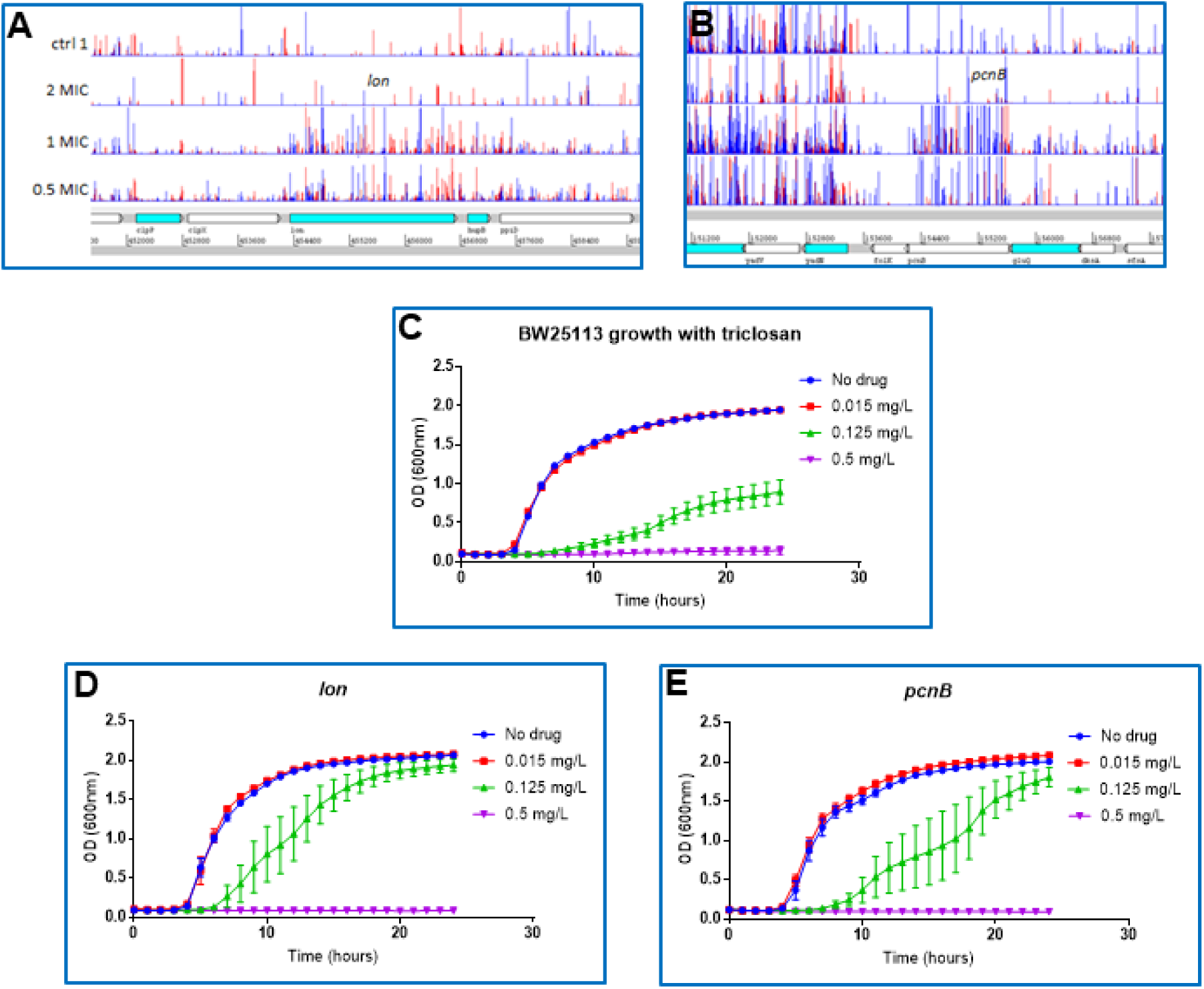
stability of mRNA and protein half-life is important to survival of exposure to triclosan. **Panels A** and **B** show insert patterns indicating inactivation of *lon* and *pcnB* is beneficial for survival in the presence of triclosan. The format of these genetic maps is as described in Figure 1. **Panels C-E** show growth curves for BW25113 and isogenic mutants grown in LB broth supplemented with different concentrations of triclosan. Growth kinetics data show mean optical density values from multiple replicates (n = 34 for BW25113 and n = 4 for other mutants). Error bars indicate standard error from the mean.

### Modulating expression of triclosan sensitivity genes confirmed predictions about the impacts of specific genes on survival in the presence of triclosan

A key advantage of using promoters in transposon mutant libraries is the ability to assay essential genes which cannot be inactivated for roles in stress responses. TraDIS+ identified numerous genes where the pattern of transposon inserts suggested expression changes were altering triclosan sensitivity. For some of these genes a growth advantage with triclosan was predicted due to transposon insertions downstream of the gene and producing an antisense transcript which will interfere with translation of mRNA. To verify these predictions, a selection of these genes were chosen for artificial up- or down-regulation. A series of constructs were made in the pBAD gene expression vector (**Figure 5**), some inserted in reverse orientation to mimic observations from the TraDIS+ data. None of the *E. coli* derivatives harbouring these plasmid constructs showed any change in growth rate on triclosan-free media. After induction with arabinose, constructs over-expressing *fabI*, *infB* and *marA* showed consistent ‘rescue’ of the ability to grow in the presence of 0.5 mg/L triclosan on agar. Constructs over-expressing *fabA*, or antisense to *rpoN* and *pstA* showed rescue of growth ability in the presence of triclosan in broth although interestingly not in agar. Artificial over-expression of *fabZ* appeared to be consistently lethal. Vector only controls, or a clone containing *lacA* (chosen as a random gene not implicated in triclosan sensitivity) showed no phenotype as expected (**Figure 5**).

**Figure 5.**
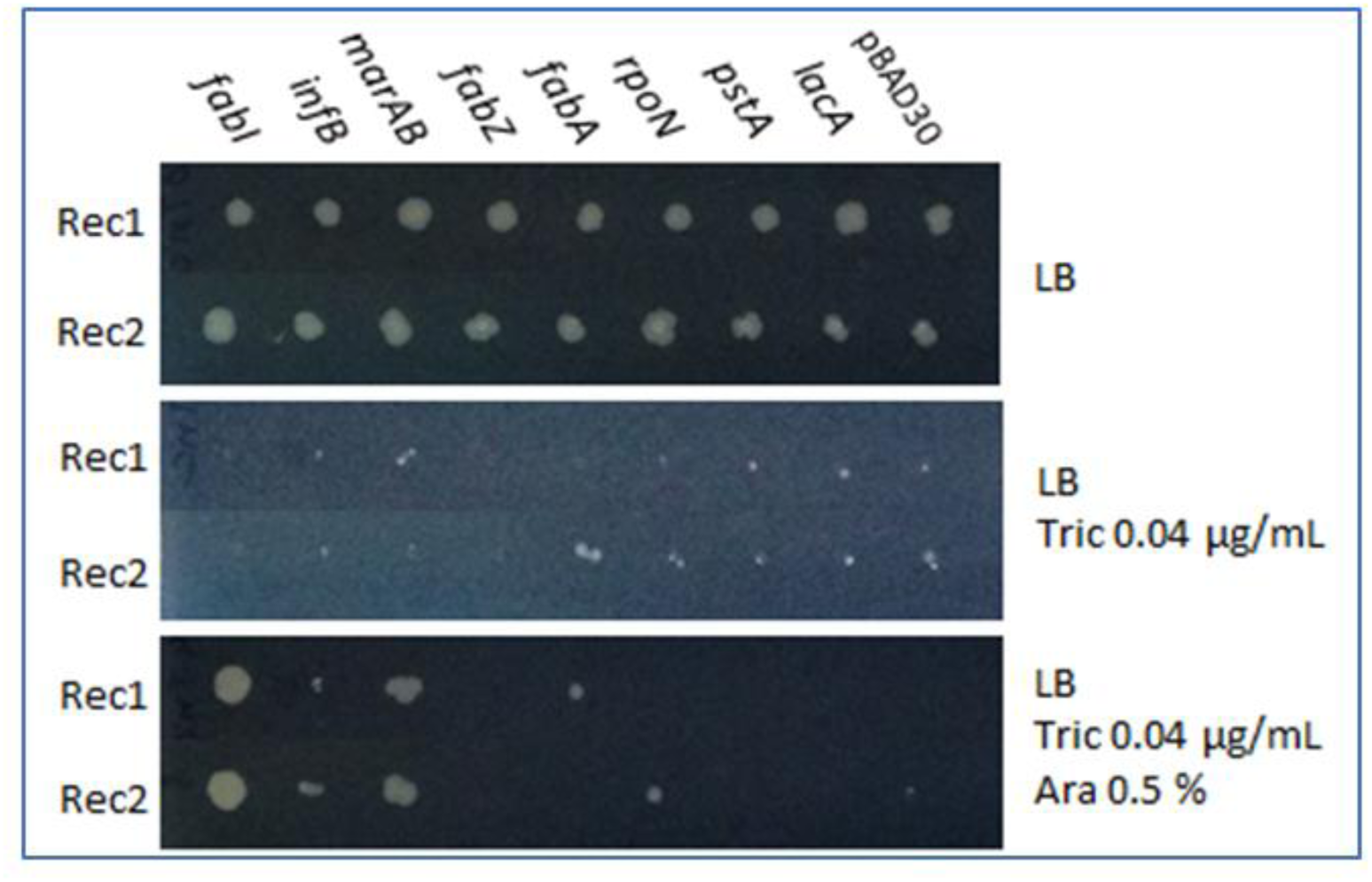
Induction or repression of genes confirms predicted roles in triclosan resistance. The coding sequences for a selection of genes were cloned into the pBAD30 expression vector under transcriptional control of the arabinose-inducible promoter. This was to modify their expression and test the predicted impact on triclosan sensitivity. Genes *fabI*, *infB*, *marAB* and *fabZ* were cloned in the forward orientation allowing their up-regulation after induction. In contrast, *fabA*, *rpoN*, *pstA* and *lacA* were cloned in the reverse direction with respect to the orientation of the *ara* promoter allowing inducible production of an antisense transcript of these genes and consequent repression of expression. Increased resistance to triclosan was predicted by up-regulation of the first four of these recombinants, and for *fabA*, *rpoN* and *pstA* by repression. The *lacA* and pBAD30 vector alone constructs were included as negative controls. Duplicate recombinants (Rec1, Rec2) for each gene were tested by spotting cultures onto LB-agar (top pane), LB agar supplemented with 0.04 ug/mL triclosan (middle pane), or LB agar supplemented with 0.04 μg/mL triclosan and 0.5 % arabinose. All recombinants grew without selection, and none grew in the presence of triclosan without arabinose. With triclosan and arabinose induction, the *fabI*, *infB* and *marAB* recombinants consistently grew better than the controls. The *fabZ* recombinant did not grow with any arabinose induction; even in the absence of triclosan (not shown), suggesting that over-expression of this gene is lethal.

## Discussion

The use of large transposon libraries and massively-parallel sequencing to assay the fitness of large numbers of mutants has proved hugely powerful in defining the essential set of genes needed for any bacteria to grow in any specific condition. For example, a recent report refined the *E. coli* essential gene list in common laboratory conditions and identified essential regions within genes^20^. Studies using the original TraDIS methodology are however limited by not being able to assay essential genes and by the time taken to make mutant libraries and sequencing costs. Improved workflows including the use of outward facing promoters of different strengths have helped to address this shortcoming^21–23^.

Here we have incorporated inducible control to an outward-facing promoter integrated into the transposon sequence, providing improved control of the expression levels of adjacent genes. This approach has also improved ‘signal to noise’ and allowed comparisons of induced and uninduced conditions to identify genes where expression changes contribute to conditional survival. The use of a single, inducible promoter, rather than a suite of transposons with promoters of different strengths helps avoid loss of density due to only a subset of each library carrying each promoter. It also avoids any site-specific insertional biases which may arise from the different sequences of a suite of different transposons. Additionally, it has proved a suitable approach to identify where ‘knock-down’ of expression of a gene can influence survival which we demonstrate has identified previously unknown targets. The technology thus enables rapid and simultaneous genome wide genotype to phenotype association determinations.

In addition, we have developed the “AlbaTraDIS” analysis software suite which performs the automated identification of significant changes in mapped sequence reads which reflect the changes in the different mutant numbers between growth conditions^24^.

‘AlbaTraDIS’ identifies significant insert sites in a context and gene-agnostic manner whilst then identifying those inserts likely to inactivate genes or alter their expression. Comparisons across diverse conditions allows commonalities and differences in responses to a stress to be easily identified, allowing analysis and integration of large numbers of data sets in an efficient manner.

We have used this approach to study mechanisms of resistance to triclosan across a wide range of drug concentrations. The data revealed the known mechanisms of triclosan resistance and importantly allowed us to identify essential genes involved in triclosan sensitivity (including the known primary target) based on expression changes. These experiments validate the use of outward facing promoters in the TraDIS+ method and shows the AlbaTraDIS software suite can identify inserts efficiently and predict impacts on gene expression in an automated manner. Having an automated analysis pipeline allows multiple conditions to be analysed in conjunction and the relationships between different stress conditions to be inferred (**Figure 2**). Using this approach, we were able for the first time to unpick the differential impacts of triclosan exposure at high and low concentrations. These data are summarised in **Figure 6** showing an integrated diagram highlighting the fundamentally distinct mechanisms of triclosan action and resistance revealed by TraDIS+.

**Figure 6.**
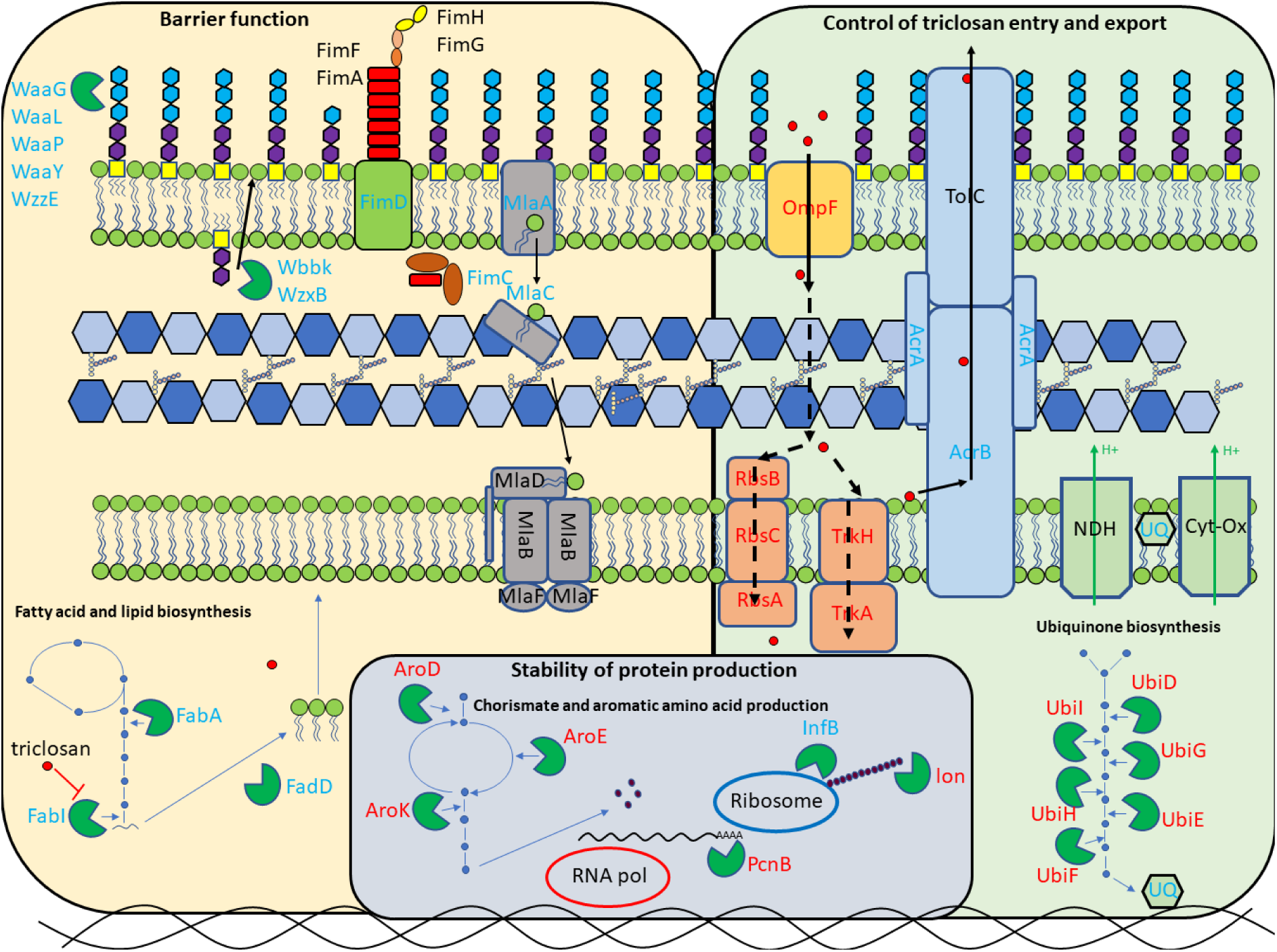
Integrated cellular diagram illustrating the major mechanisms of triclosan action and resistance. Overview of the major cellular pathways revealed as being involved in triclosan resistance by TraDIS+. Proteins with blue name labels appear beneficial to survival in the presence of triclosan, those with red name labels contribute to triclosan sensitivity. Dotted lines indicate potential routes of triclosan movement across the inner membrane.

Triclosan has become endemic in modern life and has been shown to impact microbes in the environment^7^. Our work suggests diverse residual concentrations of triclosan may result in very different selective pressures and these findings may help inform the risk of the selection of unintended cross-resistance to other drugs. There have been multiple reports suggesting links between triclosan and antibiotic resistance. A recent paper described a link between triclosan exposure and induction of levels of ppGpp which, in turn result in up to 10,000-fold levels of protection against bactericidal antibiotics^9^. Interestingly we saw no signal for *spoT* or *relA* (genes implicated in increasing ppGpp levels in that study). It may be that increasing ppGpp levels is not itself protective against triclosan, or that ‘hard-wired’ mutations are not needed to allow a fully protective response to occur after exposure and hence we observed no altered pattern of inserts at these loci in the presence of triclosan.

Multidrug efflux pumps are known to influence triclosan sensitivity. Interestingly, in our study the structural components of the pumps themselves were not protected by exposure to sub-inhibitory concentrations of triclosan although de-repression of *marA* which controls efflux expression was selected. At concentrations above the MIC, the AcrAB efflux system becomes critically important with mutants being enriched that will result in pump over-expression (through induction of SoxS or inactivation of *acrR*) as well as evident protection of *acrAB* itself. Similarly, inactivation of *mprA* with consequent augmented transcription of the *emrAB* efflux system was also observed when exposed to concentrations of triclosan above the MIC. This is important as selection of mutants that over-express these regulators is a well-known route to multidrug-resistance and suggests exposure to triclosan concentrations above the MIC is likely to select for antibiotic cross resistance.

One currently unclear aspect of triclosan’s activity relates to how it enters the cell. Whilst it is well established that triclosan crosses the outer membrane through porins, there is no current evidence about how triclosan crosses the inner membrane. Recent debate has highlighted the importance of transporters in the uptake of many small molecules rather than lipid diffusion^25^. We identified two uptake systems (for ribose and potassium) whose inactivation results in increased resistance to triclosan. Specific mutants in these systems showed increased ability to grow in the presence of triclosan (**Figure 3**) suggesting they may have a role in drug import, further work will be needed to substantiate the role of these channels in triclosan uptake.

The mechanism by which triclosan exerts a bactericidal effect is also currently elusive. We observed a pattern of genes involved in protection of mRNA and protein stability as well as translation initiation as being involved in triclosan sensitivity when exposed to high concentrations of the drug (**Figures 4 and 6**). These observations suggest the cells central information processing mechanism is compromised by high concentrations of triclosan It remains to be seen if there is one or multiple secondary targets for triclosan which confer the bactericidal effect at high concentrations.

Here we demonstrate an important improvement in transposon based functional genomics, reveal multiple new aspects of the mechanisms of action and resistance of the canonical fatty acid biosynthesis inhibitor triclosan and use this information to assess how differential exposure conditions impose distinct selective pressures. The use of an inducible promoter allows the impacts of over-expression and repression of all genes on phenotypes to be assayed efficiently. The ability to compare different conditions rapidly, and assay essential genes opens the door to high-throughput, genome scale examination of bacterial responses to various stressors. This approach can be applied to any question where the evaluation of bacterial fitness is important.

## Methods

### TraDIS+; a refined approach allowing essential genes to be assayed for roles in conditional survival

Whilst the original TraDIS protocol has proved to be hugely powerful in assaying the role of the majority of genes of any given bacteria in surviving any given stress, essential genes cannot be assayed due to the lethality of any insertions within them. To allow essential genes to be assayed for roles in any given stress, we included the use of an inducible, outward facing promoter into the transposon cassette. This allows controlled induction of transcription from the transposon insert site: inserts immediately up or downstream of an essential gene will either up- or down-regulate the gene. Changes in abundance of these inserts can therefore be assayed by our new TraDIS+ protocol. To enable this approach, we included an outward oriented IPTG-inducible *tac* promoter at the end of the kanamycin cassette, which allows controlled induction of expression with IPTG in *E. coli* (**Supplementary figure 3** shows controlled expression from the *tac* promoter in the transposon cassette as assayed using a β-galactosidase assay). We used *E. coli* strain BW25113 as a model organism for these experiments as *E. coli* has been well studied for triclosan mechanisms of action and resistance, is compatible with the use of the *tac* promoter for conditional changes in gene expression and is the parent strain for the defined KEIO mutant library, allowing access to independent inactivation mutants to validate TraDIS predictions. **Supplementary table 2** shows all strains used in this study.

### Preparation of transposomes

Transposon Tnp001 is a mini-Tn*5* transposon coding for kanamycin resistance (*aph*(*3’*)*-Ia*) which incorporates an outward oriented, IPTG-inducible *tac* promoter, 3’ to the kanamycin resistance gene. Transposon DNA was amplified by PCR using oligonucleotides (primers are in **Supplementary table 3**), and transposomes were prepared by mixing purified transposon DNA with EZ-Tn5 transposase according to the supplier’s instructions (Lucigen Corp). The mixture was incubated at 37°C for 1 h then stored at −20°C before use.

### Preparation of a BW25113 mutant library

*E. coli* strain BW25113 was prepared for electroporation as previously described^26^. Bacteria were grown in 2X YT broth to an optical density of 0.3 (measured at 600 nm). After harvesting, bacteria were washed three times in 10% glycerol before finally being suspended in 10% glycerol. Electrotransformations were performed using a Bio-Rad ‘Gene Pulser II’ electroporator set to 2400 V and 25 μF, coupled to a ‘Pulse Controller II’ set to 200 Ω and using 2 mm electrode gap cuvettes. Immediately after the electric pulse, cells were suspended in 1 ml of super optimal broth with catabolite repression (S.O.C. medium) (NEB) and incubated at 37°C for 1 h. The cell suspension was then spread on LB agar supplemented with kanamycin at 30 mg/L and incubated overnight at 37°C. Resulting colonies were harvested and pooled together in 15% glycerol in LB broth for storage at −70C.

### Triclosan exposure conditions

To determine the exposure conditions for experiments, the minimum concentration of triclosan needed to inhibit growth of BW25113 (MIC) was determined using a 2-fold dilution method in LB broth, the same growth medium that was used to perform the TraDIS+ experiments. In addition, viable counts of BW25113 were performed to determine bactericidal/bacteriostatic impacts by preparing duplicate two-fold serial dilutions of triclosan from 4 to 0.008 mg/L in LB broth, inoculated with approximately 10^6^ cfu/ml and incubated at 37°C. Cell numbers at 0 h were obtained from a 10 µl sample, removed immediately from each culture and serially diluted 10-fold to a dilution of 10^−7^ in a 96-well plate. 5µl volumes of the dilutions were then spotted onto LB agar, and the spots allowed to dry prior to incubation overnight at 37°C. After the LB broth cultures plus triclosan dilutions had been incubated at 37°C for 24 h, a second 10 µl sample was removed from each and treated similarly to obtain viable counts after 24 h in the presence of triclosan.

For TraDIS+ experiments, approximately 10^7^ mutants from the transposon mutant library pool were grown in LB broth supplemented with a range of concentrations of triclosan in doubling increments ranging from 0.008 mg/L to 1 mg/L representing a 125-fold range of concentrations. Experiments were also completed in the absence of IPTG or in the presence of either 0.2 or 1 mM IPTG to induce transcription from the transposon outward oriented promoter. Control experiments were also performed in the presence of two ethanol concentrations (ethanol was used to dissolve triclosan). All experiments were performed in duplicate. A total of 66 independent TraDIS+ experiments were completed.

### Preparation of customised sequencing library

The pooled mutant mixtures were grown overnight in LB broth in the various conditions outlined above, DNA was extracted using a ‘Quick-DNA™’ Fungal/Bacterial 96 kit extraction kit (Zymo Research). A Nextera DNA library preparation kit (Illumina) was used to prepare DNA fragments for nucleotide sequencing except that Tnp-i5 oligonucleotides were used instead of i5 index primers, and 28 PCR cycles were employed. The amplified DNA fragment mixture was selected for a size range of 300bp-500bp and sequenced on a NEXTSeq 500 sequencing machine using a NextSeq 500/550 High Output v2 kit (75 cycles). Negative control PCRs were included from BW25113 DNA and from mutant library DNA which had not been subjected to tagmentation to confirm specificity of amplification.

### Bioinformatics

Sequence reads from Fastq files generated from TraDIS sequencing, were mapped to the BW25113 reference genome (CP009273,^24^) and analysed using the Bio-TraDIS (version 1.4.1,^18^) and AlbaTraDIS (version 0.0.5, ^24^) toolkits. This process includes filtering out of any reads without a transposon tag sequence. Bio-TraDIS was used to align sequence reads (using SMALT version 0.7.4) to the reference genome and to create insertion plots. To identify gene essentiality for the different conditions, genes that have very few or no insertions are identified. Within the Bio-TraDIS toolkit (tradis_essentiality.R), a threshold value for the number of insertions within essential genes is estimated using the observed bimodal distribution of insertion sites over genes when normalized for gene length^19, 27^. To compare the insertion patterns between different concentrations of Triclosan (conditions) and in media only (control), the AlbaTraDIS pipeline was used. As the TraDIS+ approach also provides data on the induction and repression of genes, we were interested in changes in frequency of insertion patterns, not only within genes, but also in the regions flanking the gene. A preferential increase of insertions oriented towards the gene (5’ to 3’) in the upstream region is suggestive of modulation of expression. A preferential increase of insertions in the antisense direction of the coding region of a gene (3’ to 5’) in the downstream region is suggestive of inhibition of gene expression by production of an interfering RNA transcript. To identify these patterns, the AlbaTraDIS toolkit (albaTraDIS script using -a flag for use of annotation) calculates the number of ‘forward’ and ‘reverse’ insertions per gene and 198 bp upstream and downstream of the gene (using annotation for CP009273,^28^) for all genes in all conditions and control. Then the number of sequence reads are modelled on a per-gene basis using a negative binomial distribution and an adapted exact test as implemented in edgeR^29^ followed by multiple testing correction^30^ is used to identify significant differences (using default cut-offs of q-value <= 0.05, logFC >= 1, logCPM > 8) in insertion frequency between each condition and control. The result is a comprehensive comparison that predicts which genes are important under the test stress as well as an indication if a change in expression of a gene (either down- or up-regulated) has certain advantages for survival in the different concentrations of triclosan.

Mapping of insert sites were visually inspected using ‘Artemis’ which was used to capture images of selected areas of the genomes of interest with insertions for figures^31^.

### Validation experiments

A total of 82 mutants were selected from the KEIO library of defined mutants to validate predictions from the TraDIS+ data set^32^. These were selected to include genes representing the major pathways identified as involved in triclosan sensitivity as well as a set of six randomly selected genes with no predicted impact on triclosan sensitivity. The KEIO library contains two, independent insertional inactivated mutants for each gene and both were used for each of the 82 targets. Two triclosan resistant mutants were also included - a BW25113 mutant selected after exposure to high level triclosan (as in ^17^) and L702, a previously characterised highly triclosan resistant mutant of *Salmonella* Typhimurium^17^. Both carried substitutions within FabI (Gly93Val).

For each mutant, the MIC of triclosan was determined as previously detailed. In addition, the ability of all mutants to grow in the presence of three concentrations of triclosan (0.015, 0.125 and 0.5 mg/L) was determined over 24 hours and compared to drug-free controls to allow for generic growth impacts to be identified. Experiments were run in 96 well format in 100 µl of LB broth inoculated with ~1000 cfu. Growth was assessed by measuring absorbance of cultures at 600nm every 60 min in a BMG FLUOStar OMEGA plate reader. Each experiment was duplicated giving four datasets for each gene (two for each mutant allele, each repeated independently). BW25113 was included in all experiments giving a total of 34 separate measurements for growth of the parent in each condition. Differences in growth were analysed by ANOVA or t-tests (when comparing single mutant vs BW25113 at one condition). Growth capacity of each mutant in all conditions was also analysed by determining velocities of logarithmic growth and by calculating the area under the curve values for the whole growth period.

The fitness of selected mutants was also determined by counting viable numbers of cells over 24 hours after exposure to triclosan concentrations as described above.

### Examining the impacts of directed alteration of gene expression on triclosan sensitivity

To confirm predicted impacts of altering expression of target genes, a selection of genes of interest were cloned into the pBAD30 vector under control of an arabinose-inducible promoter using *Eco*RI and *Xba*l restriction sites. These constructs were then tested for their ability to grow in the presence of different concentrations of triclosan on agar and in broth, both in a range of arabinose concentrations to induce expression of the cloned genes.

### Data availability

All sequence data have been deposited with EBI. The overall project accession number is PRJEB29311 and individual experiment accession numbers are:

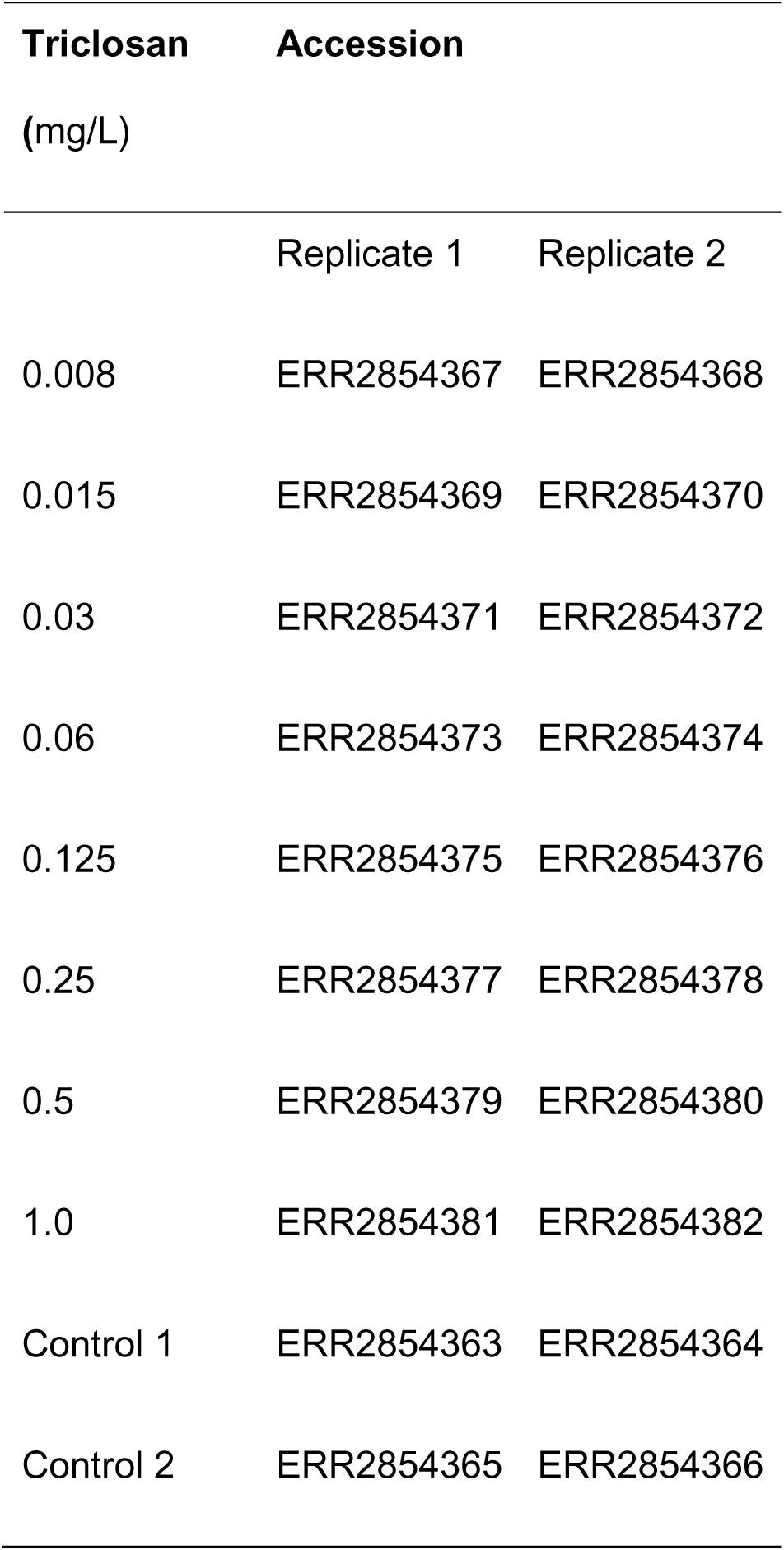

| Triclosan<br>(mg/L) | Accession |  |
| --- | --- | --- |
|  | Replicate 1 | Replicate 2 |
| 0.008 | ERR2854367 | ERR2854368 |
| 0.015 | ERR2854369 | ERR2854370 |
| 0.03 | ERR2854371 | ERR2854372 |
| 0.06 | ERR2854373 | ERR2854374 |
| 0.125 | ERR2854375 | ERR2854376 |
| 0.25 | ERR2854377 | ERR2854378 |
| 0.5 | ERR2854379 | ERR2854380 |
| 1.0 | ERR2854381 | ERR2854382 |
| Control 1 | ERR2854363 | ERR2854364 |
| Control 2 | ERR2854365 | ERR2854366 |

Growth kinetic data are provided in supplementary information

## Funding

The author(s) gratefully acknowledge the support of the Biotechnology and Biological Sciences Research Council (BBSRC); SB, AKT, MY, MAW and IGC were supported by the BBSRC Institute Strategic Programme Microbes in the Food Chain BB/R012504/1 and its constituent project BBS/E/F/000PR10349. AJP was supported by the Quadram Institute Bioscience BBSRC funded Core Capability Grant (project number BB/CCG1860/1). Genomic analysis used the MRC ‘CLIMB’ cloud computing environment supported by grant MR/L015080/1.

We would like to thank David Baker for assistance with sequencing

## Transparency declaration

The funders had no role in study design, data collection and analysis, decision to publish, or preparation of the manuscript.

## Supplementary material

### Supplementary figures

**Supplementary figure S1.**
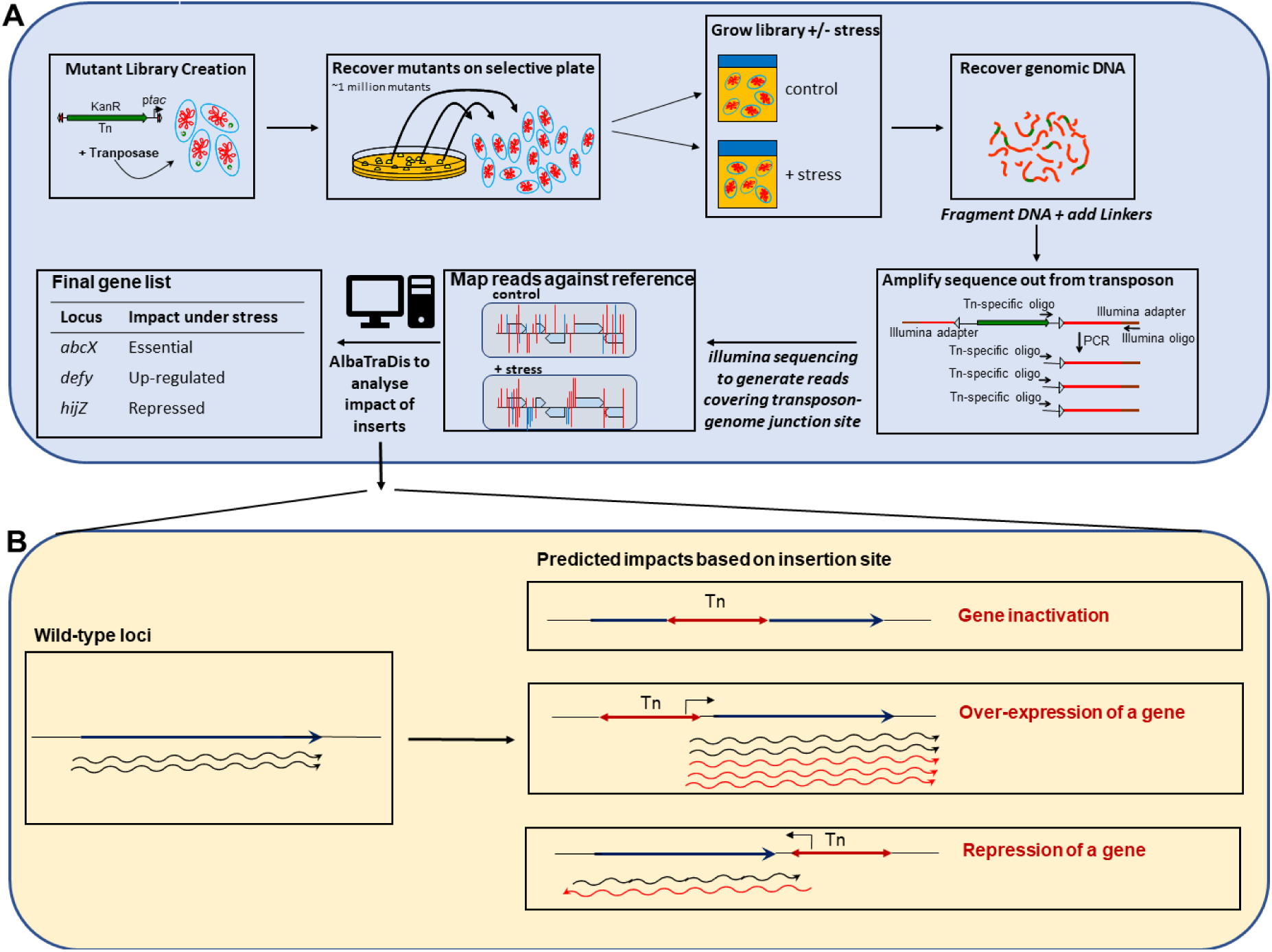
Overview of the TraDIS+ workflow. **Panel A.** Illustration of the steps in generation of mutant library, growth under stress and analysis of insert sites used in TraDIS+. **Panel B**. Example of how AlbaTraDIS predicts impacts on genes based on insert positions. Expression from the outward facing promoter is controllable at different levels using induction with IPTG.

**Supplementary Figure S2.**
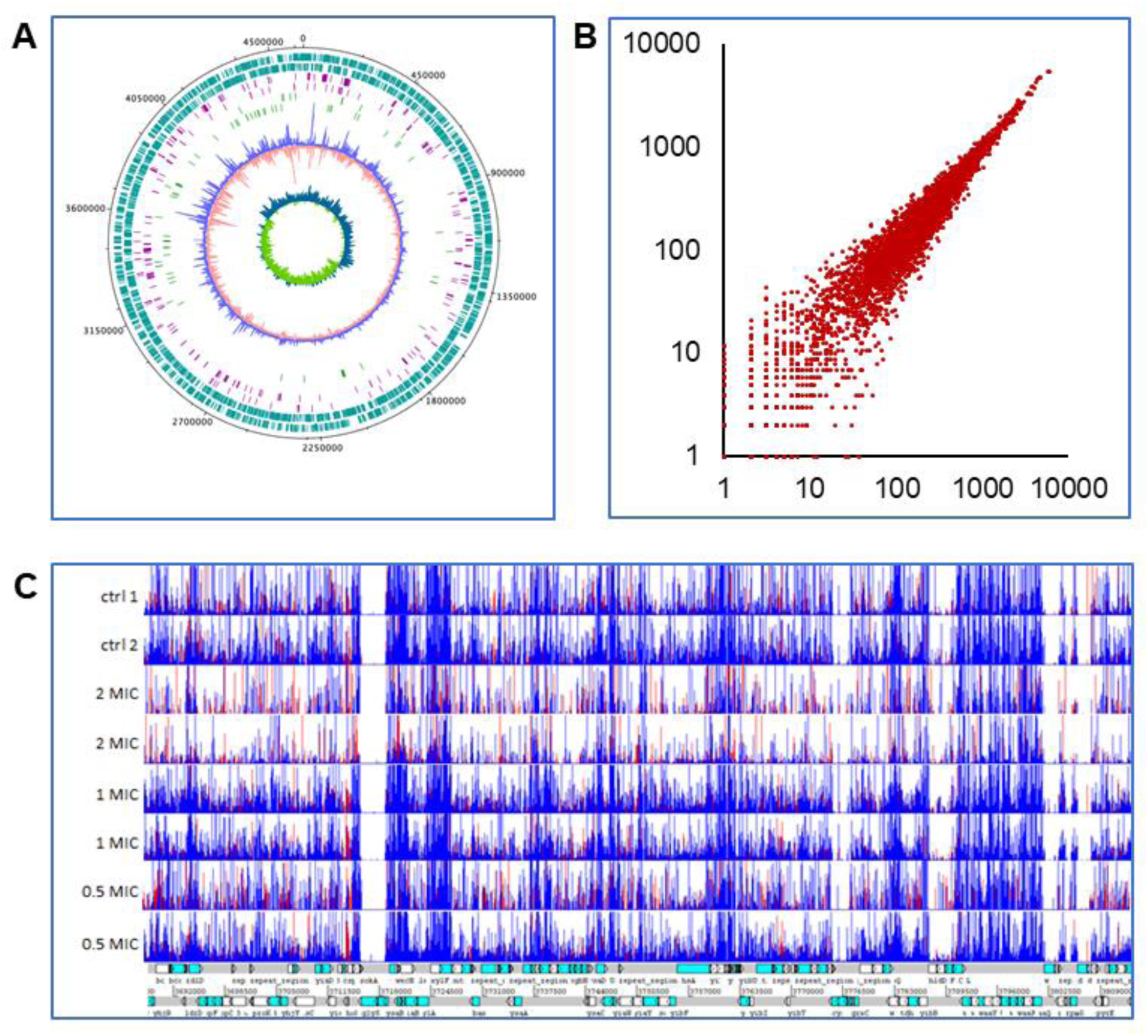
Density of the transposon library, coverage across the genome and reproducibility between replicates. **Panel A** shows density of inserts across the genome of BW25113. From outside to in: Track 1, forward genes in BW25113; Track 2, reverse genes in BW25113; Track 3, essential genes in BW25113 (‘forward strand’); Track 4, essential genes in BW25113 (‘reverse strand’); Track 5, positions of genes showing conditional role in triclosan survival (‘forward’ strand); Track 6, positions of genes showing conditional role in triclosan survival (‘reverse’ strand); Track 7, ‘forward’ insertions in control library grown in LB 0 m*M* IPTG; Track 8, ‘reverse’ insertions in control library grown in LB 0 m*M* IPTG; Track 9, GC content. Insertion density around the origin is higher due to multiple copies present because the cells were growing exponentially prior to transposon mutagenesis. **Panel B** shows a scatter plot comparing numbers of inserts isolated from two replicate libraries across the genome (points represent genes), the correlation coefficient gives an r value of 0.987 showing the consistency between replicates. **Panel C** shows a 118 kb region of the *E. coli* BW25113 genome with mapped insertion sites to demonstrate the reproducibility of the data between biological replicates. A genetic map of the region is shown along the bottom. Above this, in each of the 8 horizontal windows, vertical lines show the location of transposon insertion sites, and the height of each bar reflects the number of transposon-directed sequence reads that mapped to each site. Red bars indicate insertions are orientated in line with the forward strand and blue bars (plotted over red) orientated in line with the reverse strand. Data from separate cultures are shown for each of four exposure conditions (no triclosan controls, 2X MIC, 1X MIC and 0.5X MIC of triclosan).

**Supplementary Figure S3.**
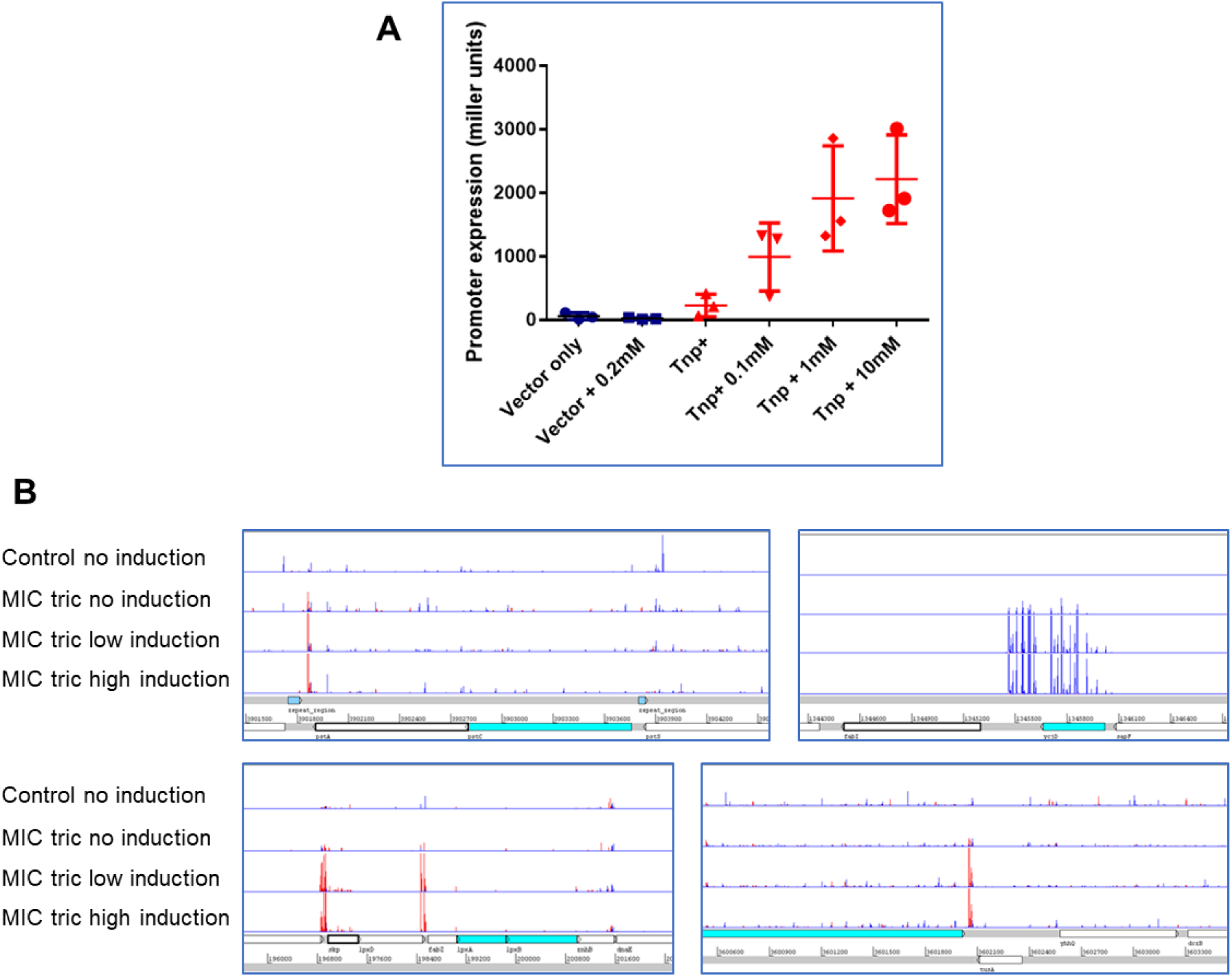
Demonstration of inducible expression from the *tac* promoter in the transposon cassette. **Panel A.** Data show miller units from β-galactosidase assays after different exposures to IPTG (concentrations marked), individual data points are shown along with mean (wide horizontal bars), and standard deviation (shorter horizontal bars). **Panel B** shows orientation specific insertions observed under transposon promoter non-inducing and inducing (+IPTG) conditions. Genetic context is shown along the bottom. Above these, in each of the 4 horizontal windows, vertical lines show the location of transposon insertion sites, and the height of each vertical line reflects the number of transposon-derived sequence reads that mapped to each site. Red lines indicate insertions are orientated in line with the forward strand and blue bars (plotted over red) orientated in line with the reverse strand. Data from independent cultures are shown for each of four exposure conditions (control with no induction, MIC of triclosan with no IPTG induction; MIC of triclosan with low IPTG induction; MIC of triclosan with high IPTG induction.

### Supplementary tables

**Supplementary table 1.**
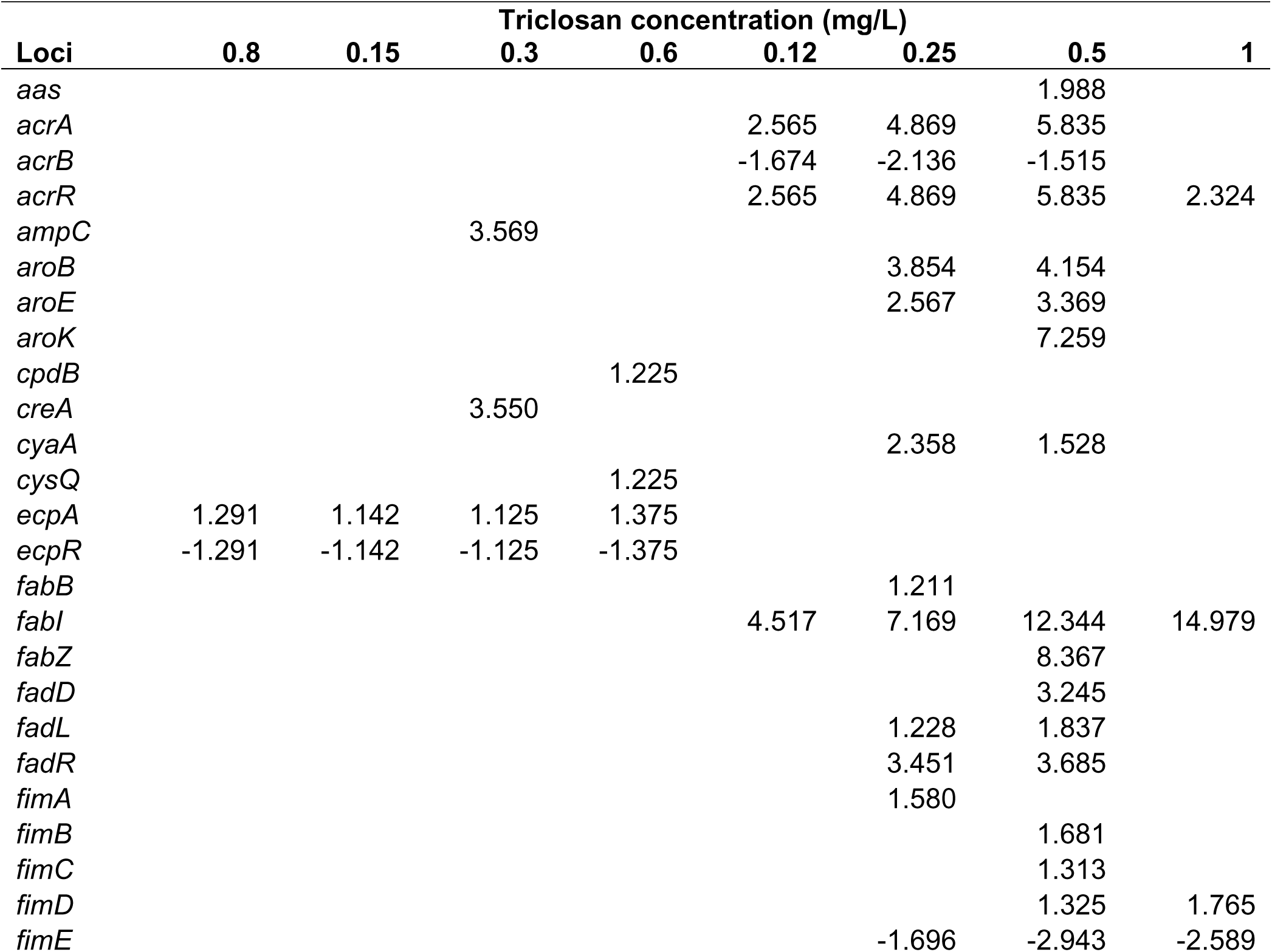
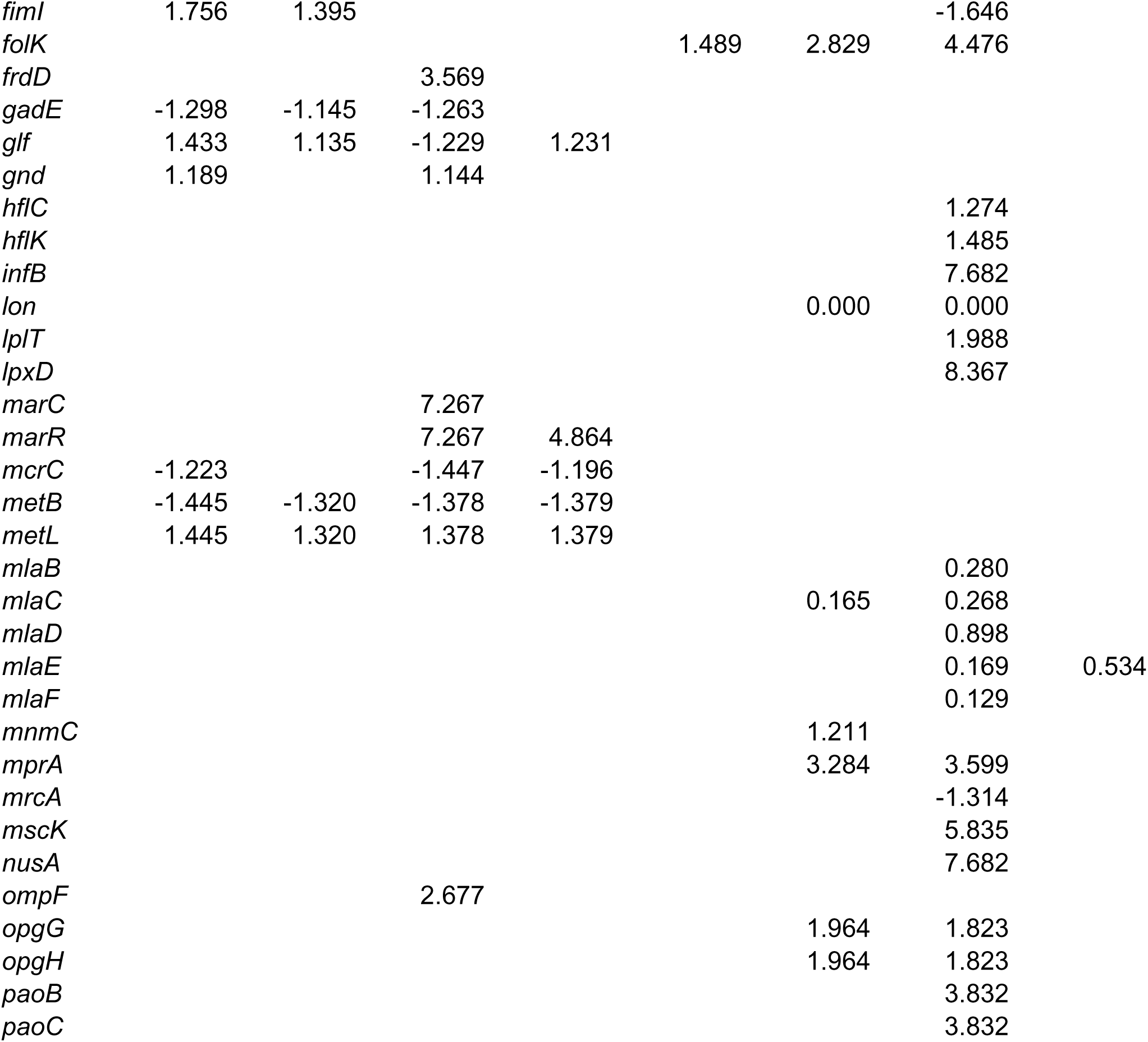
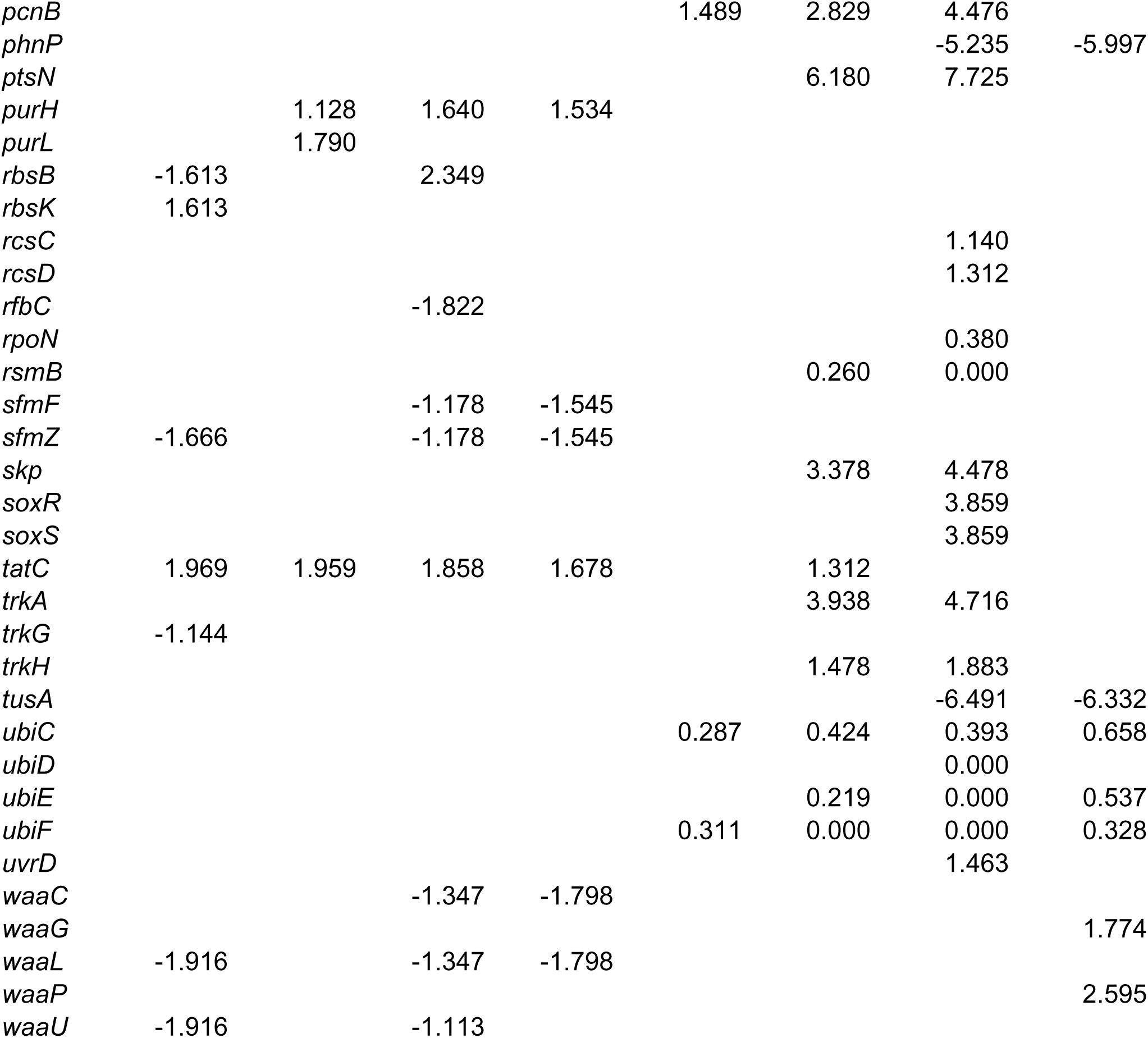
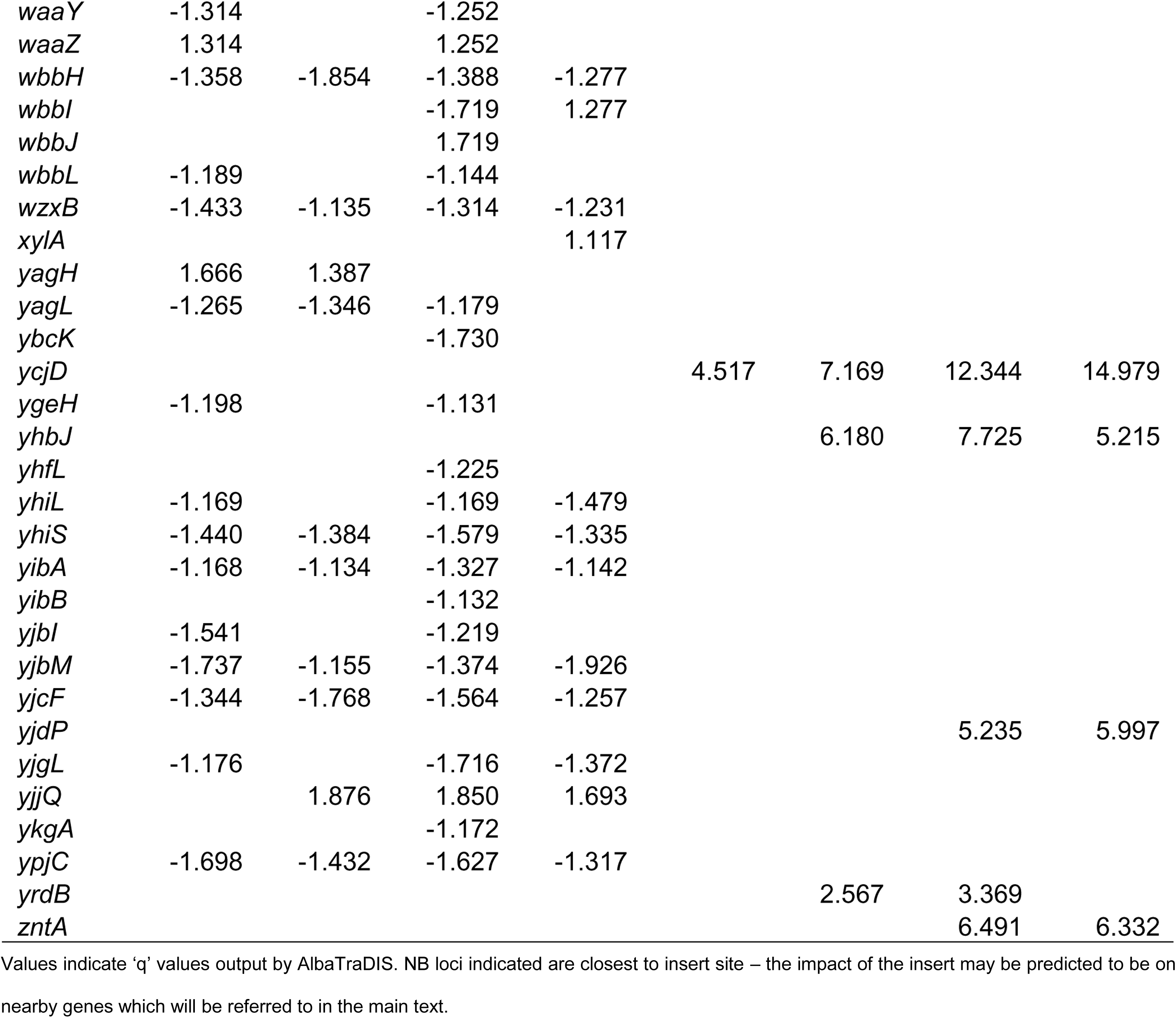
List of loci identified as significant after exposure to different triclosan concentrations

**Supplementary table 2.**
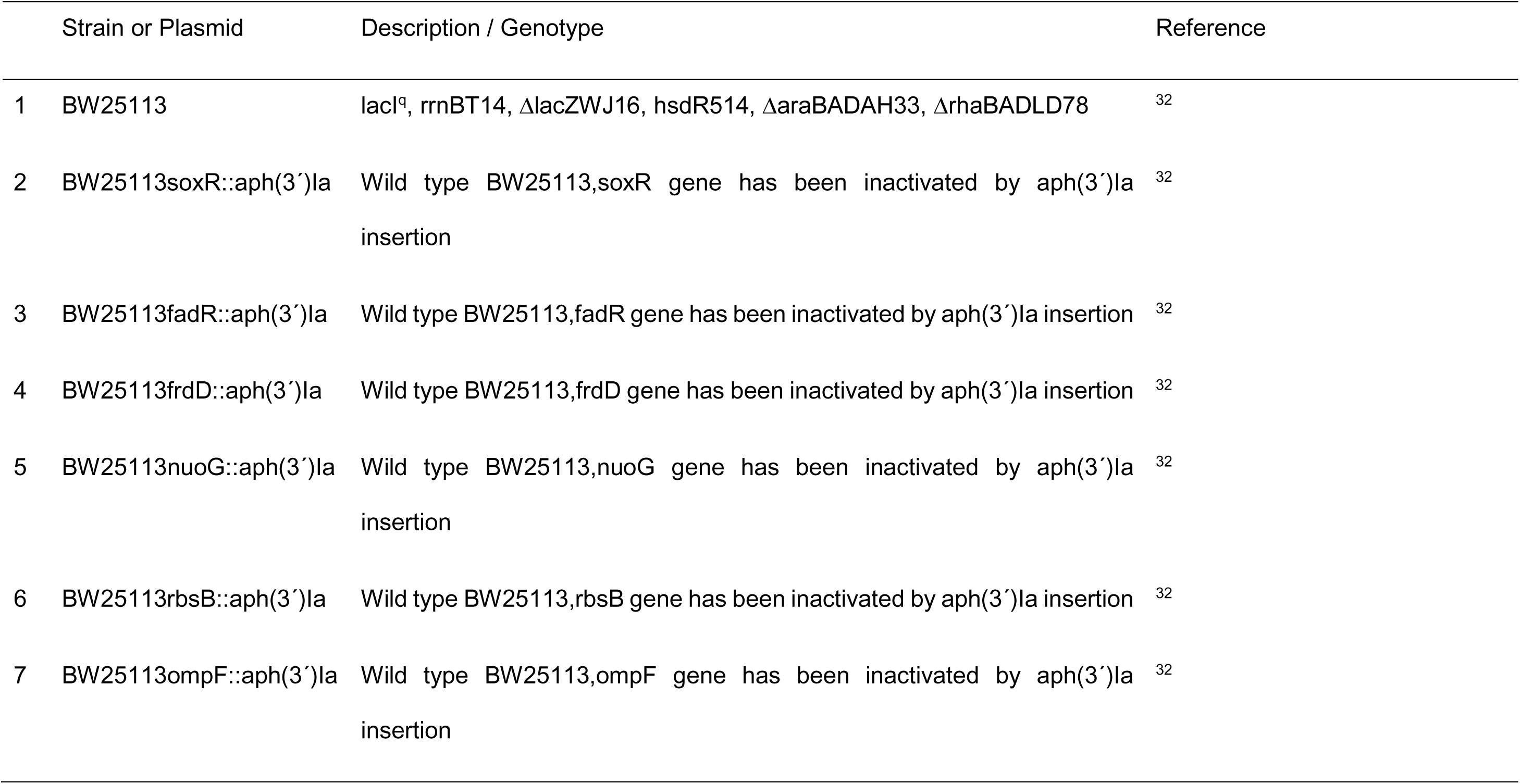
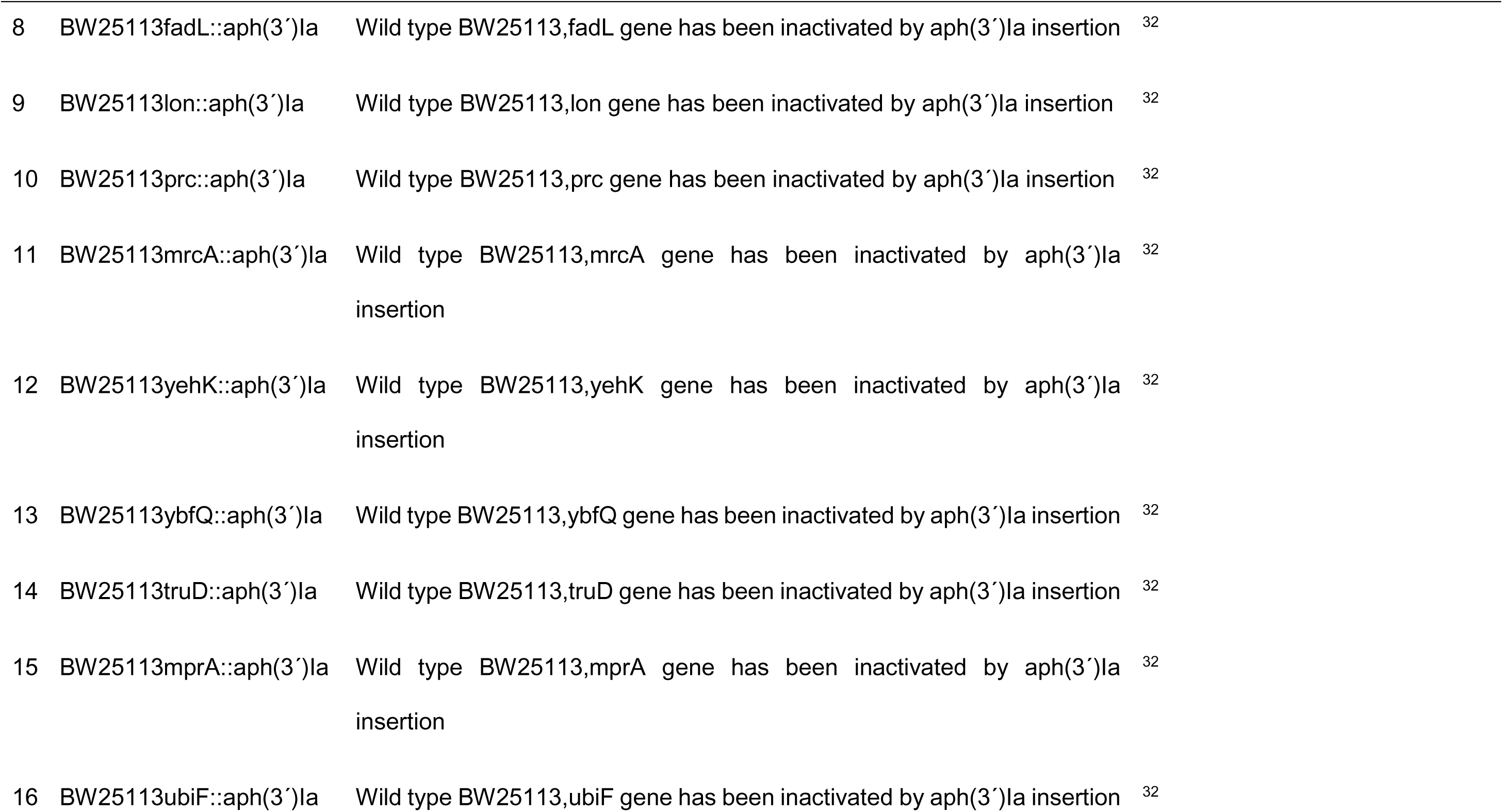
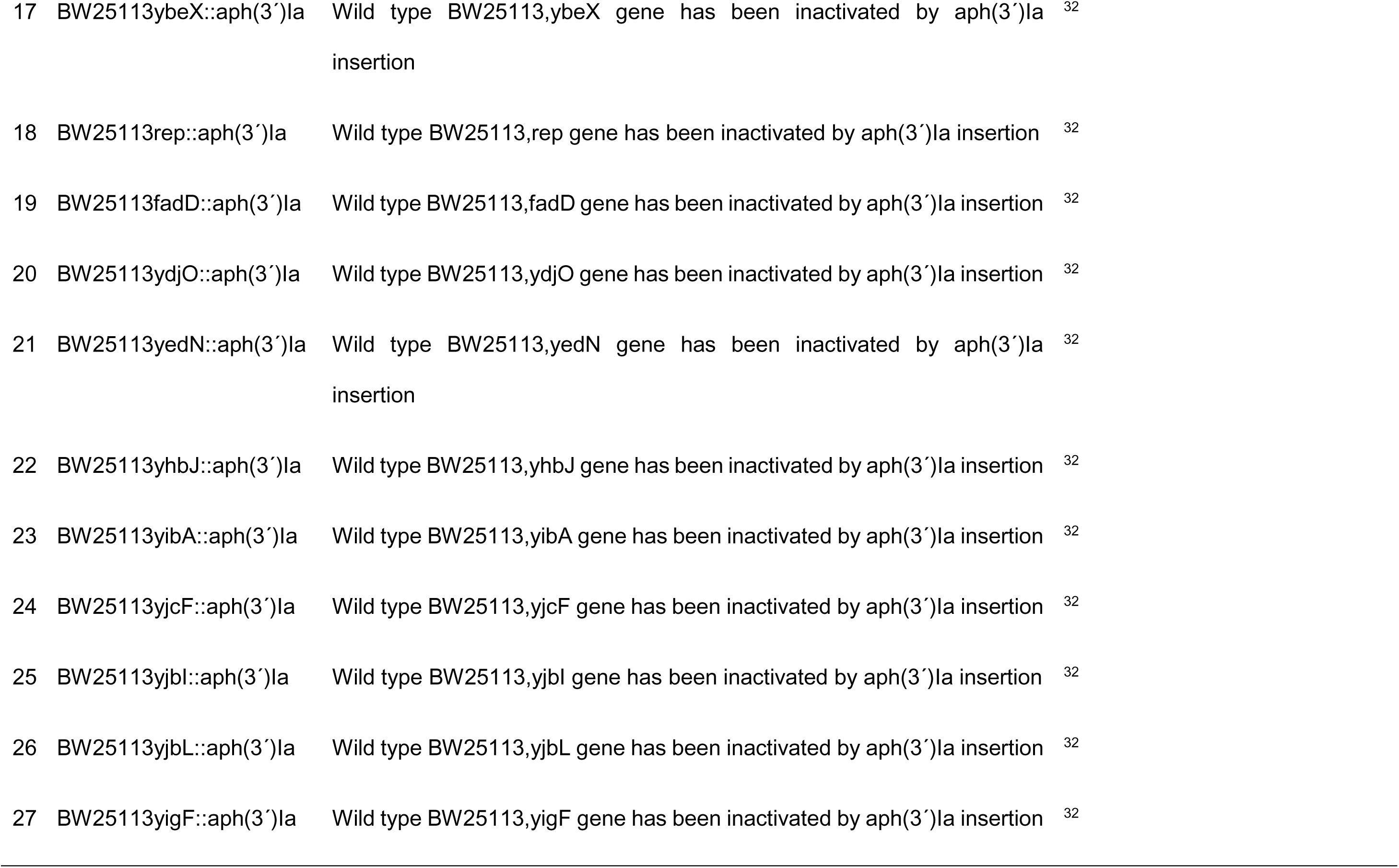
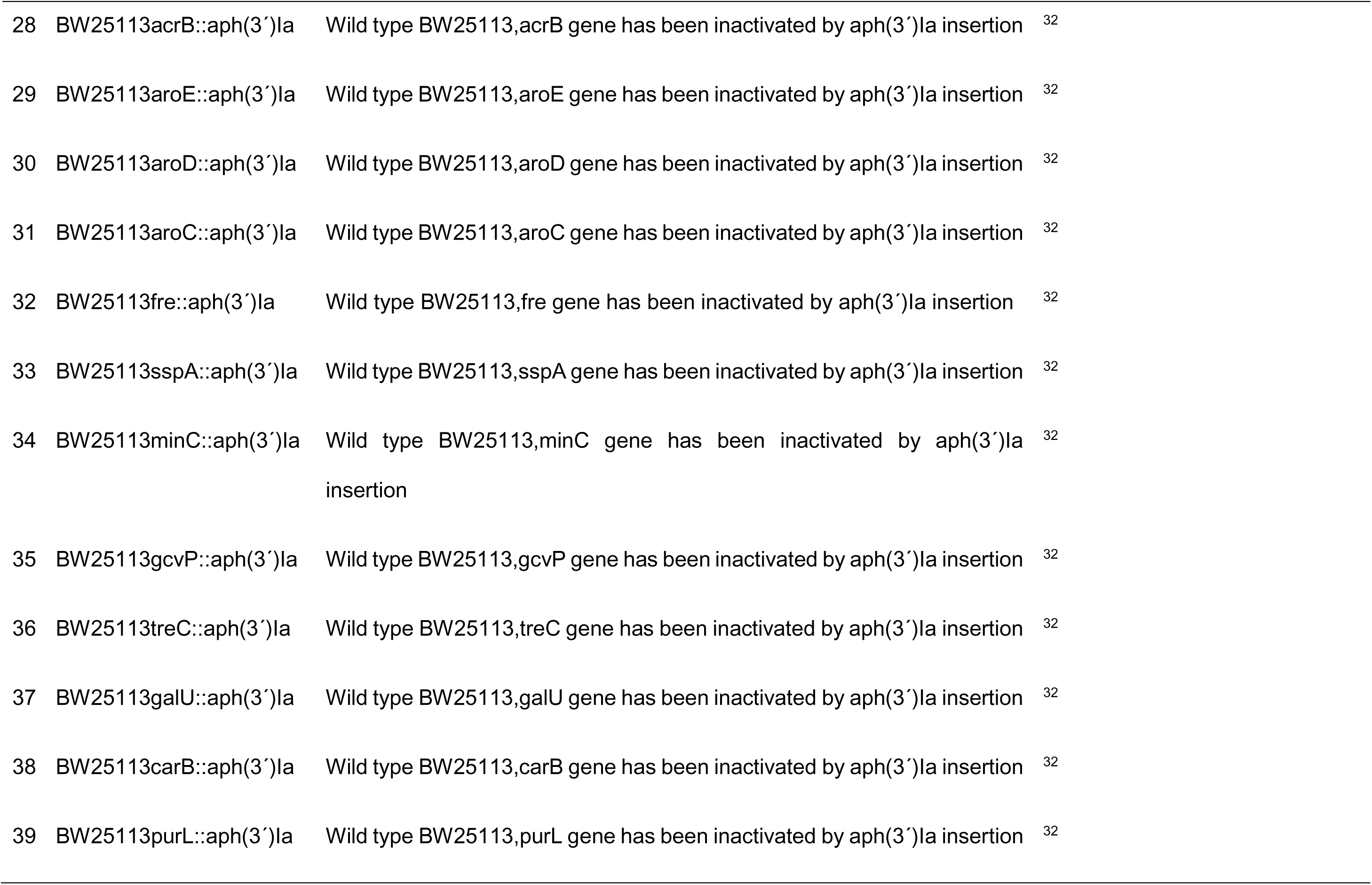
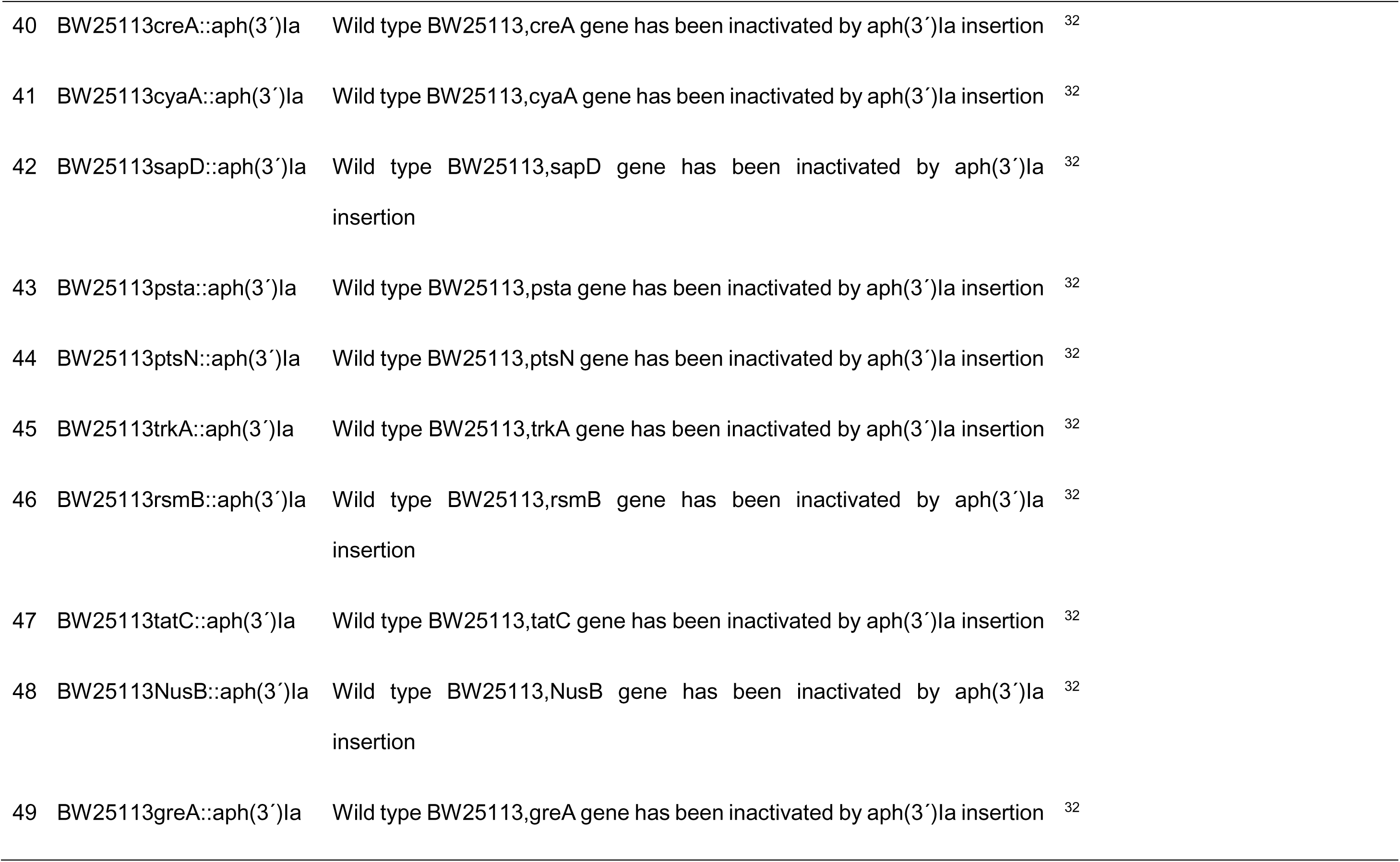
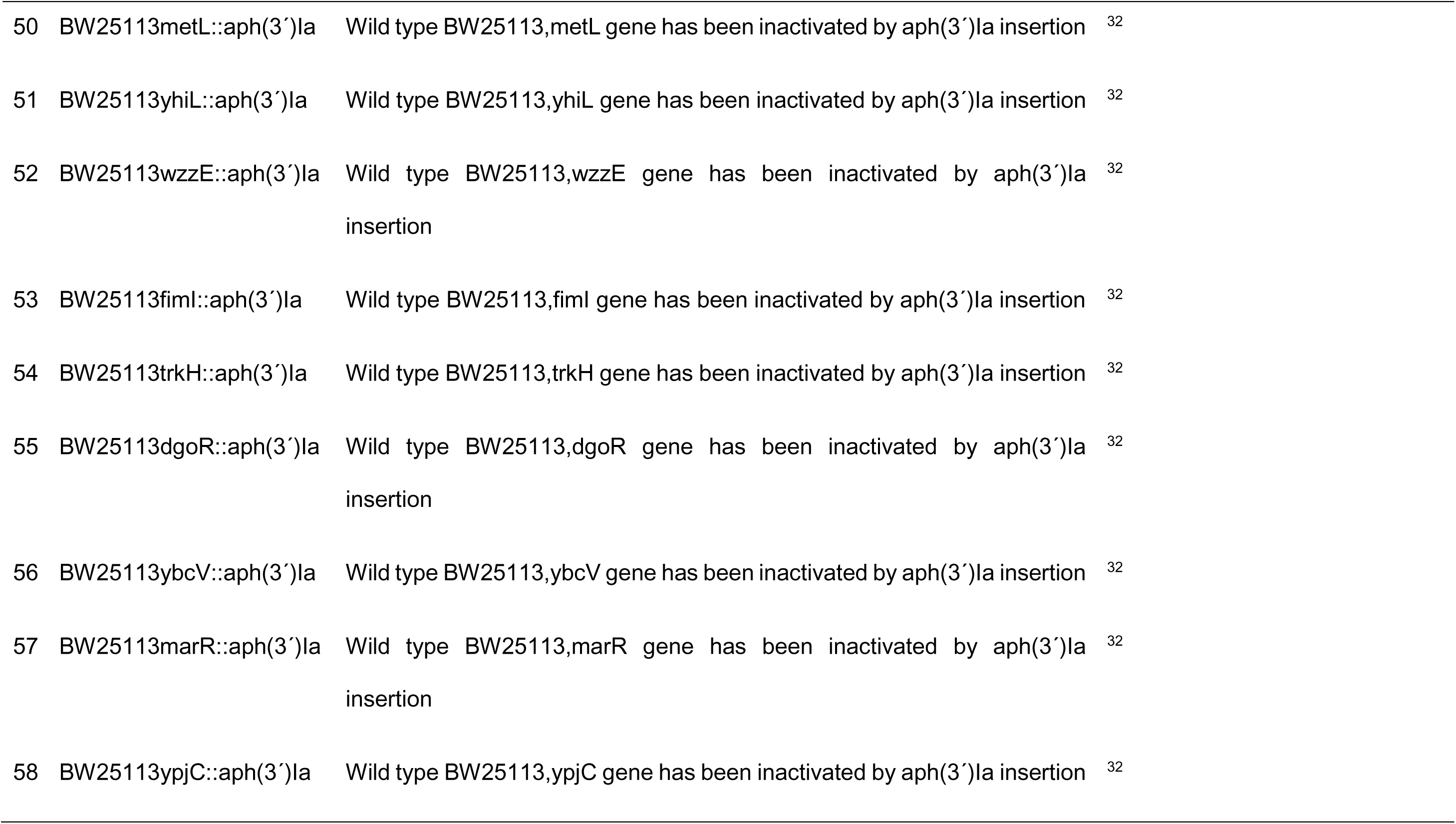
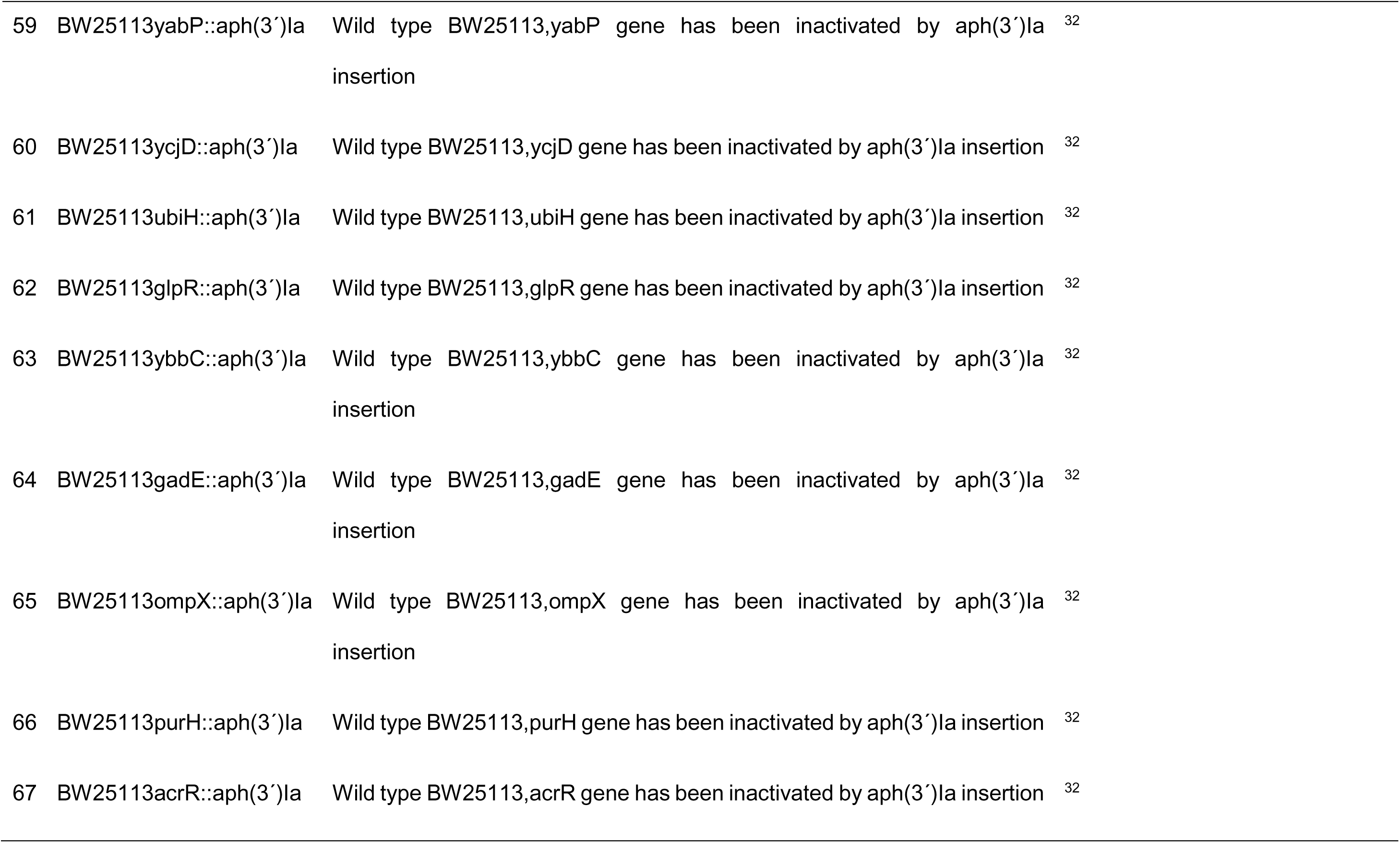
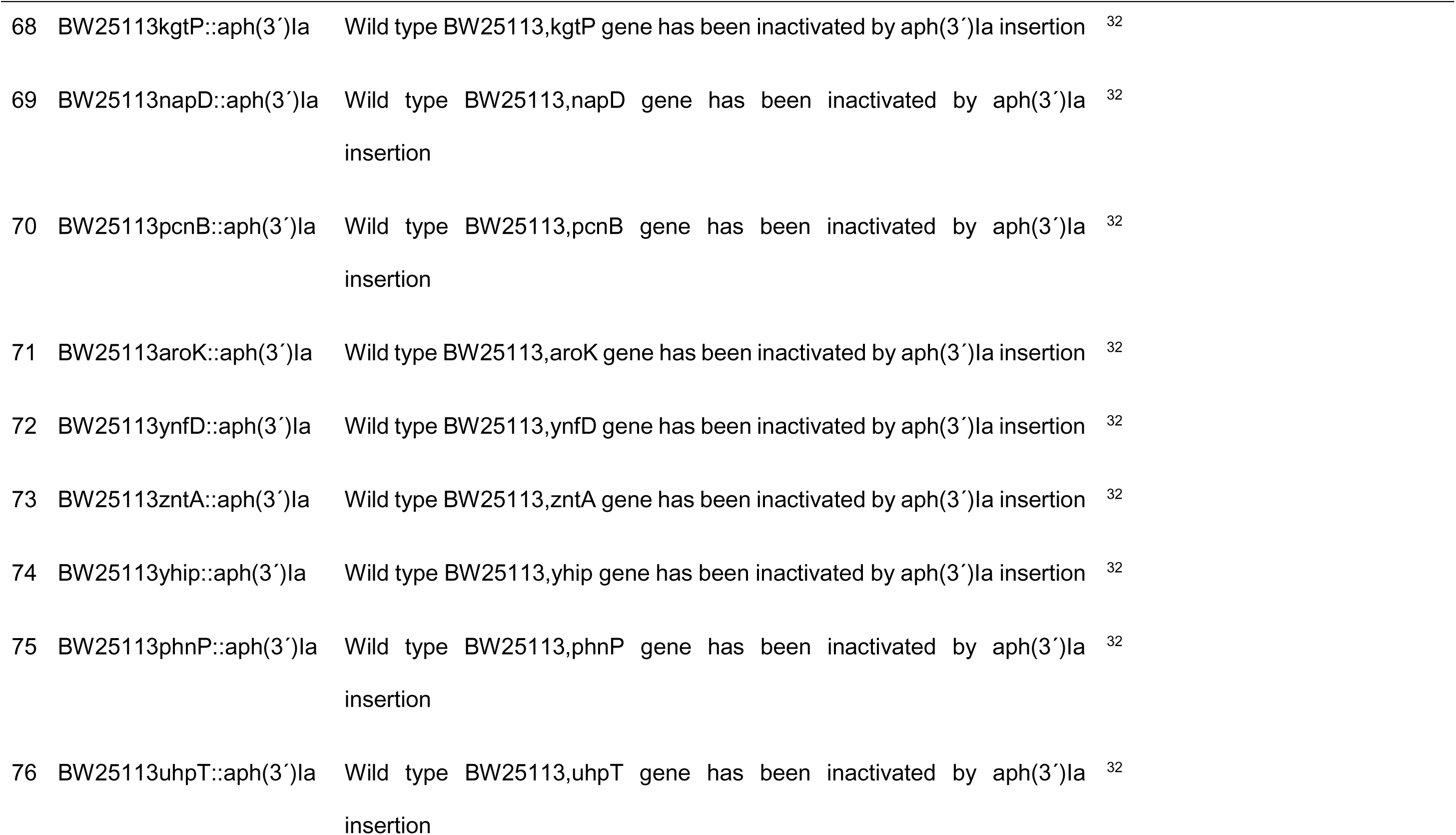
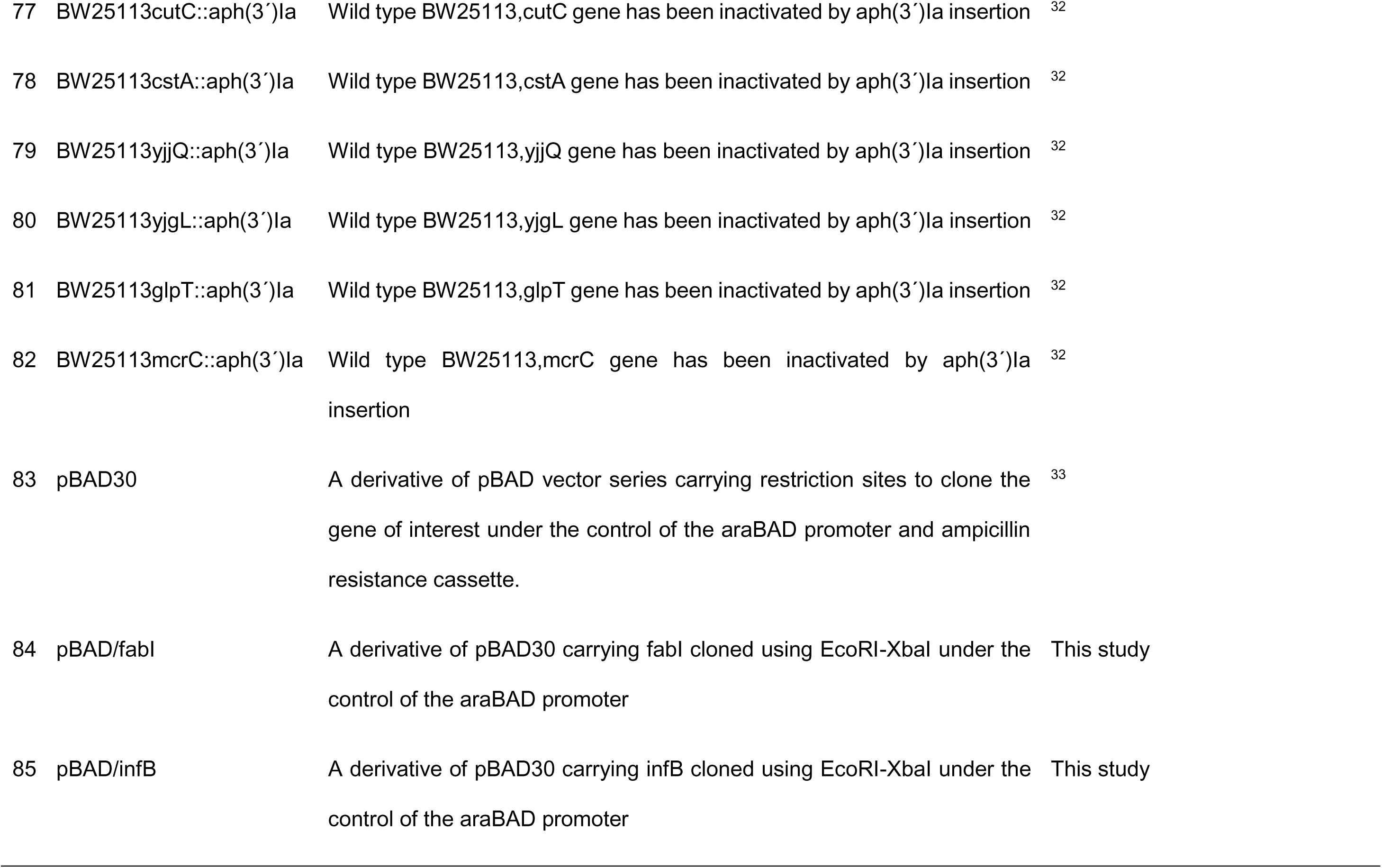
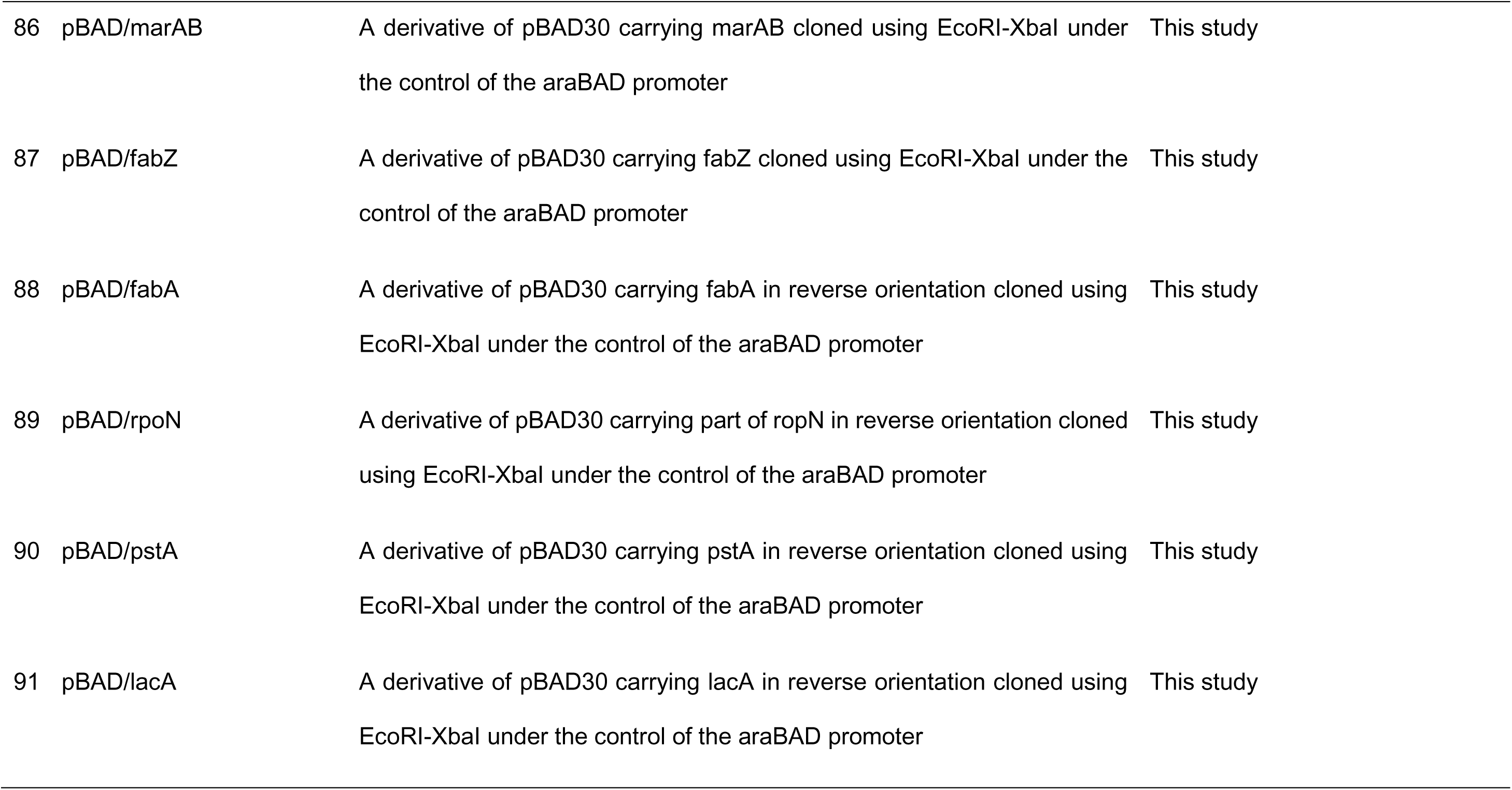
Strains, vectors and primers

**Supplementary table 3.**
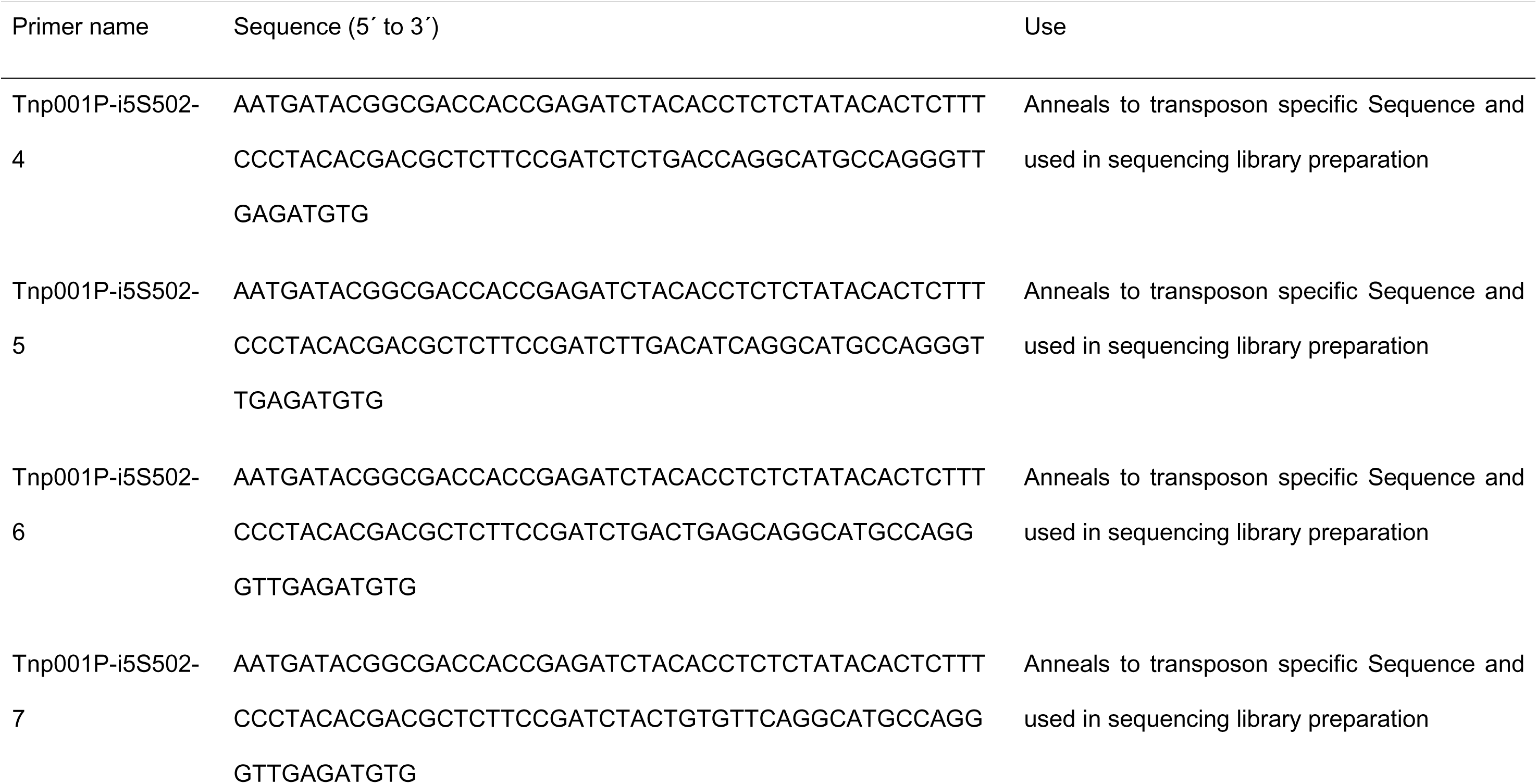
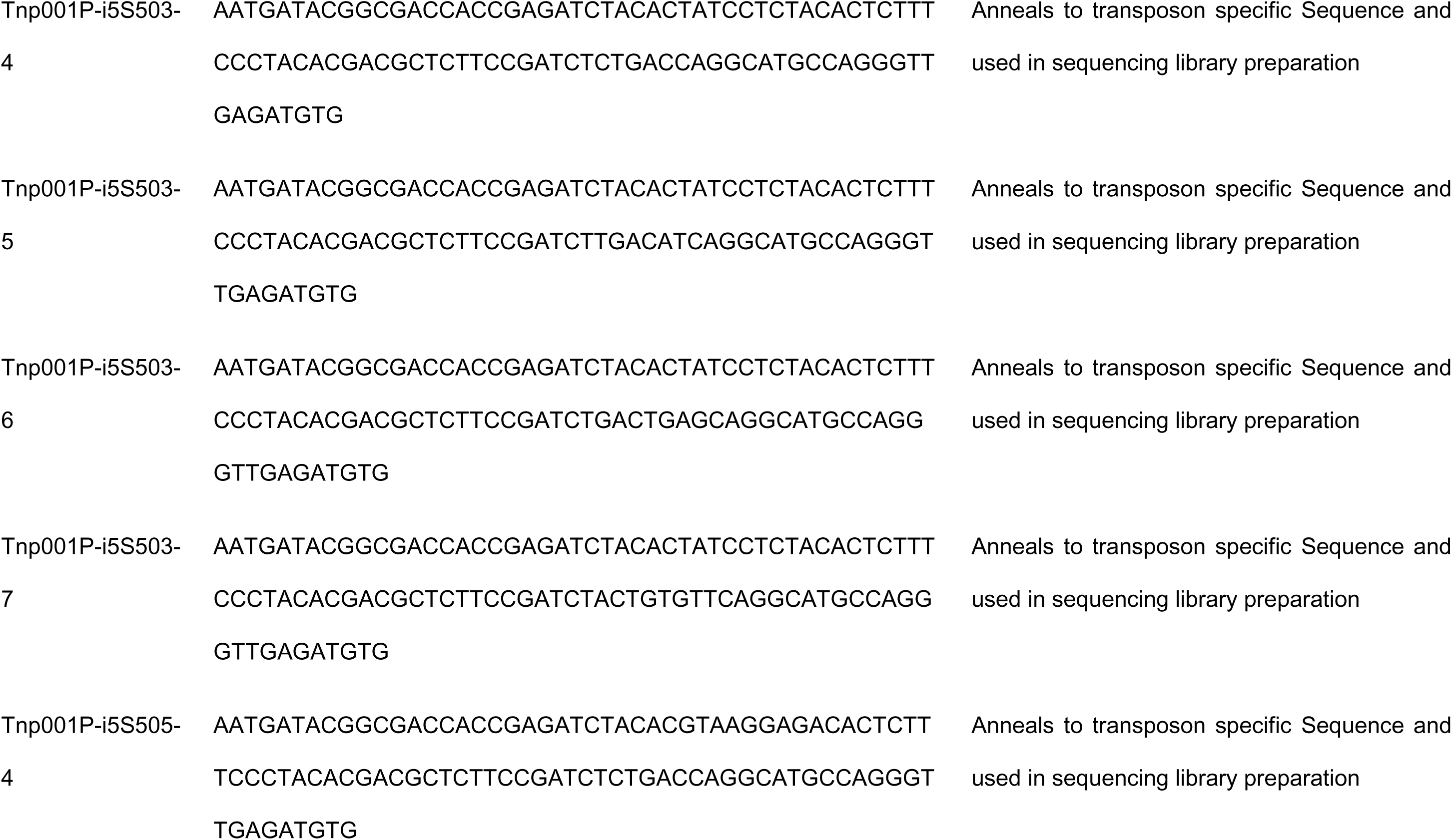
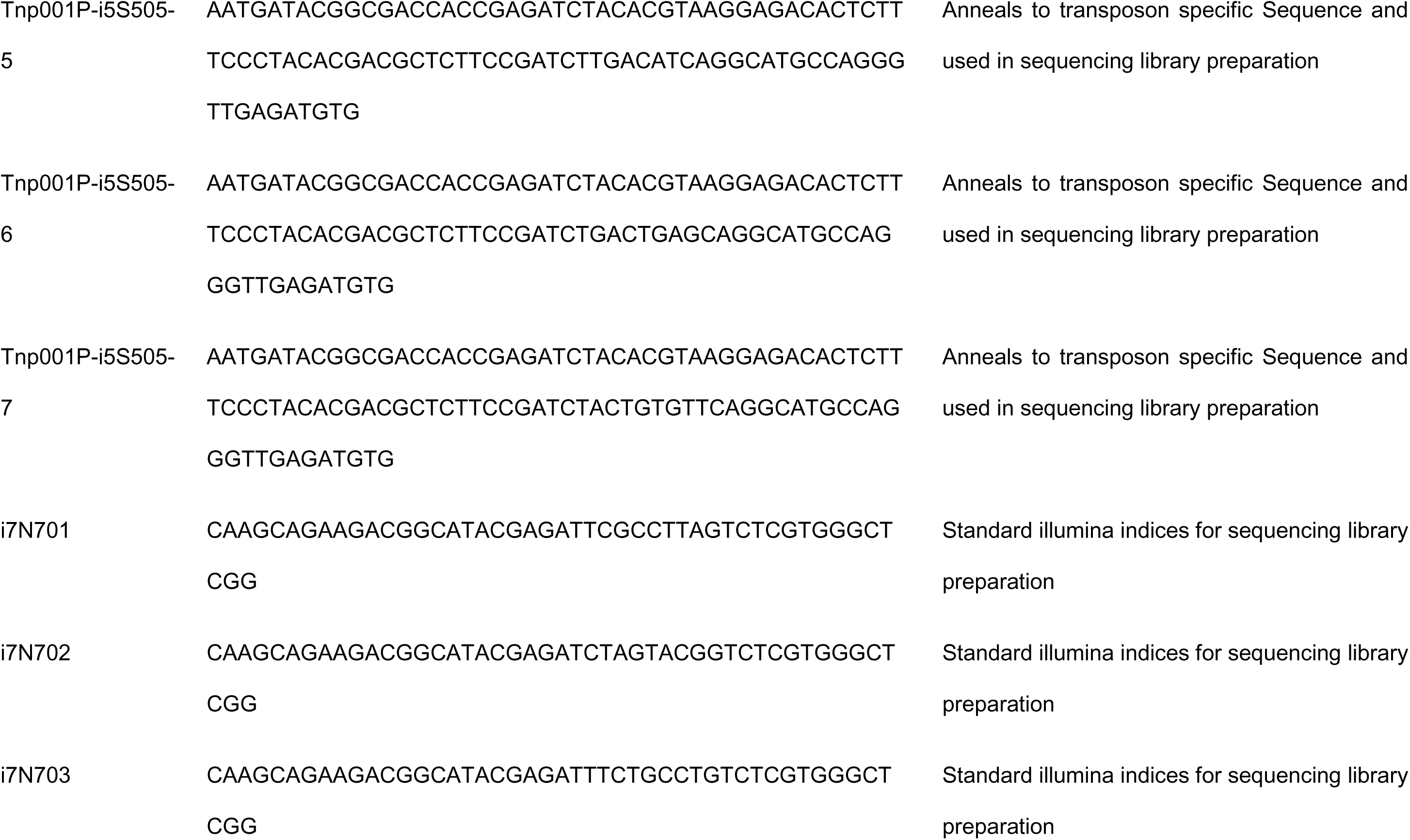
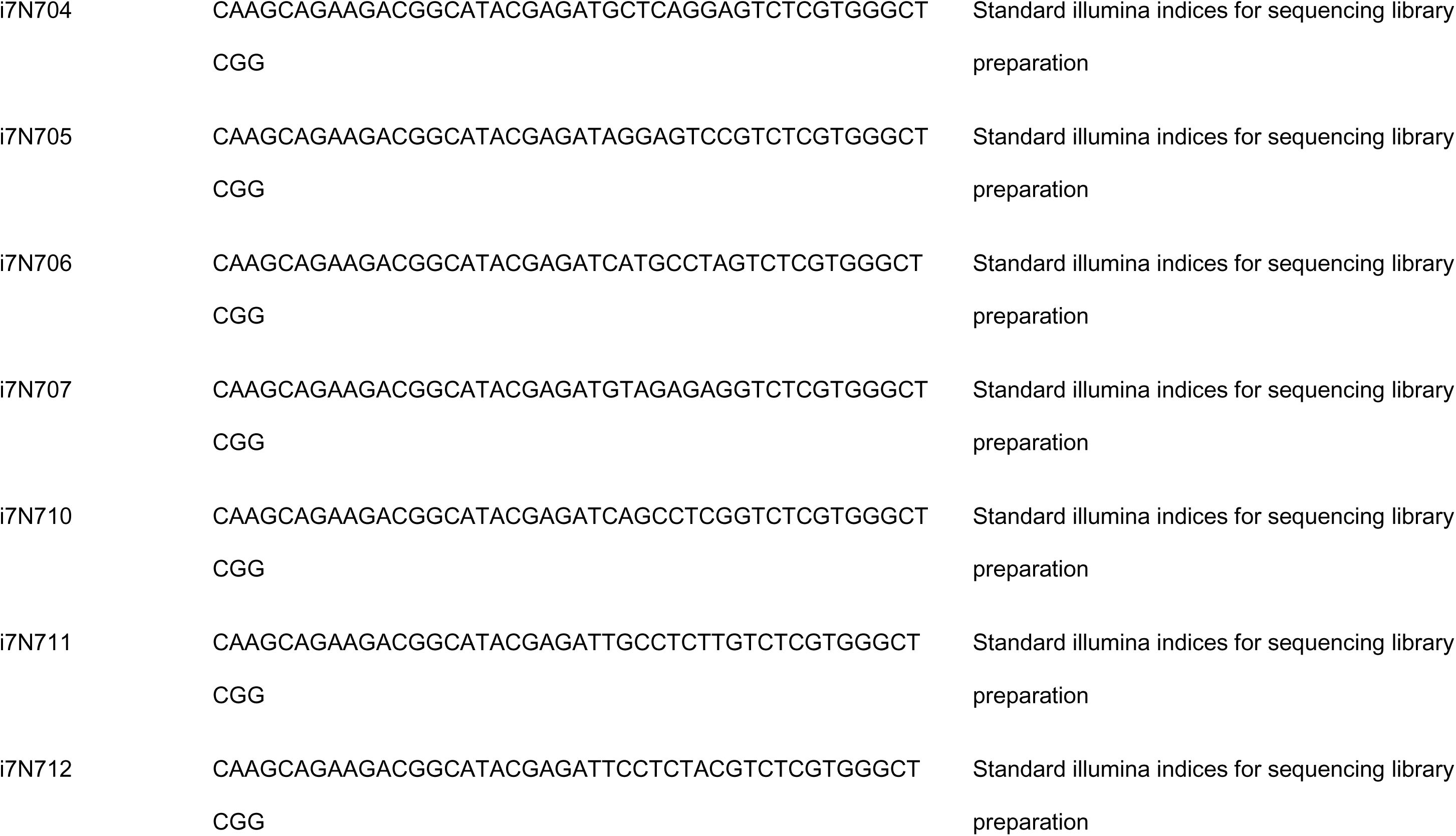
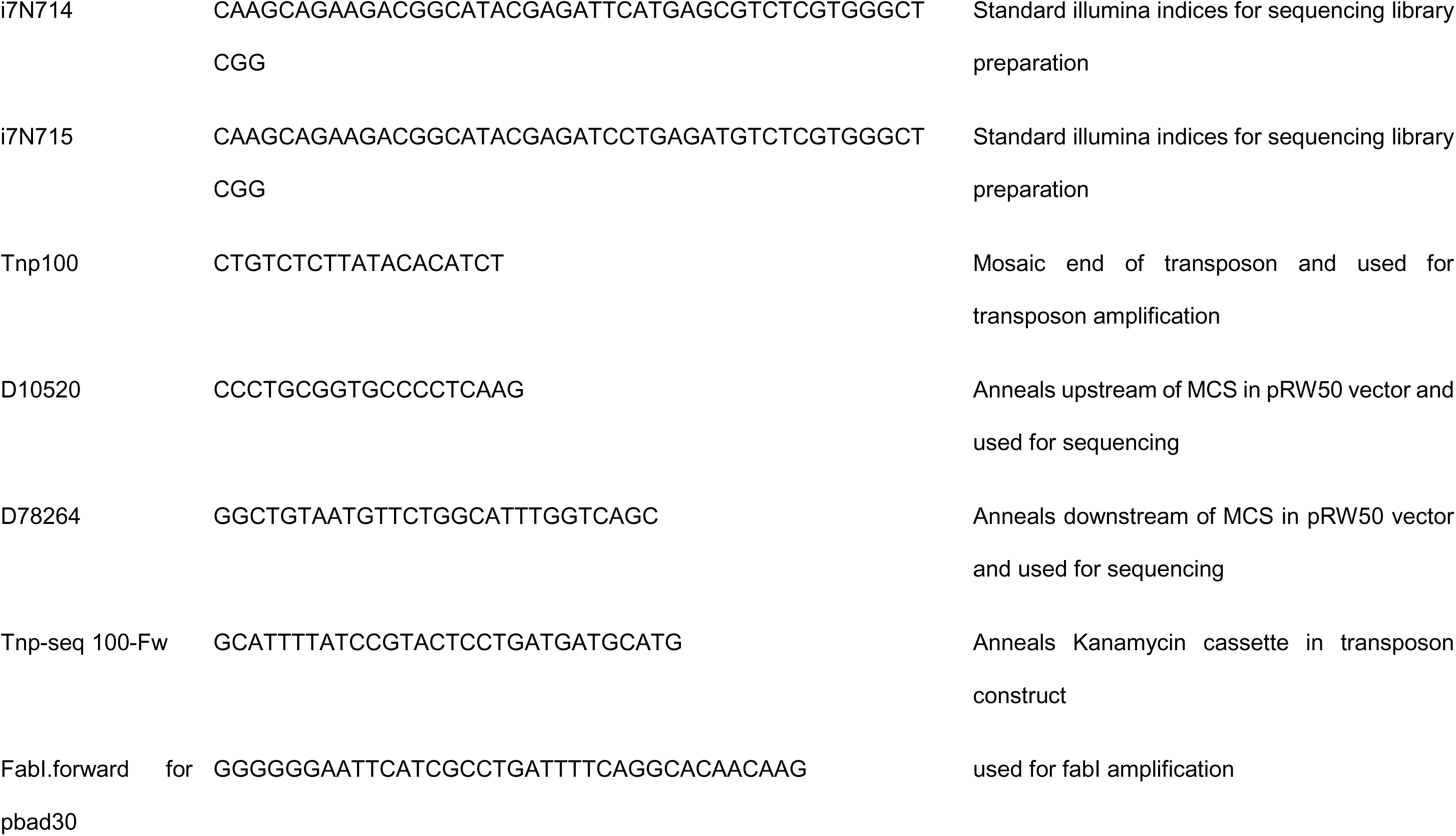
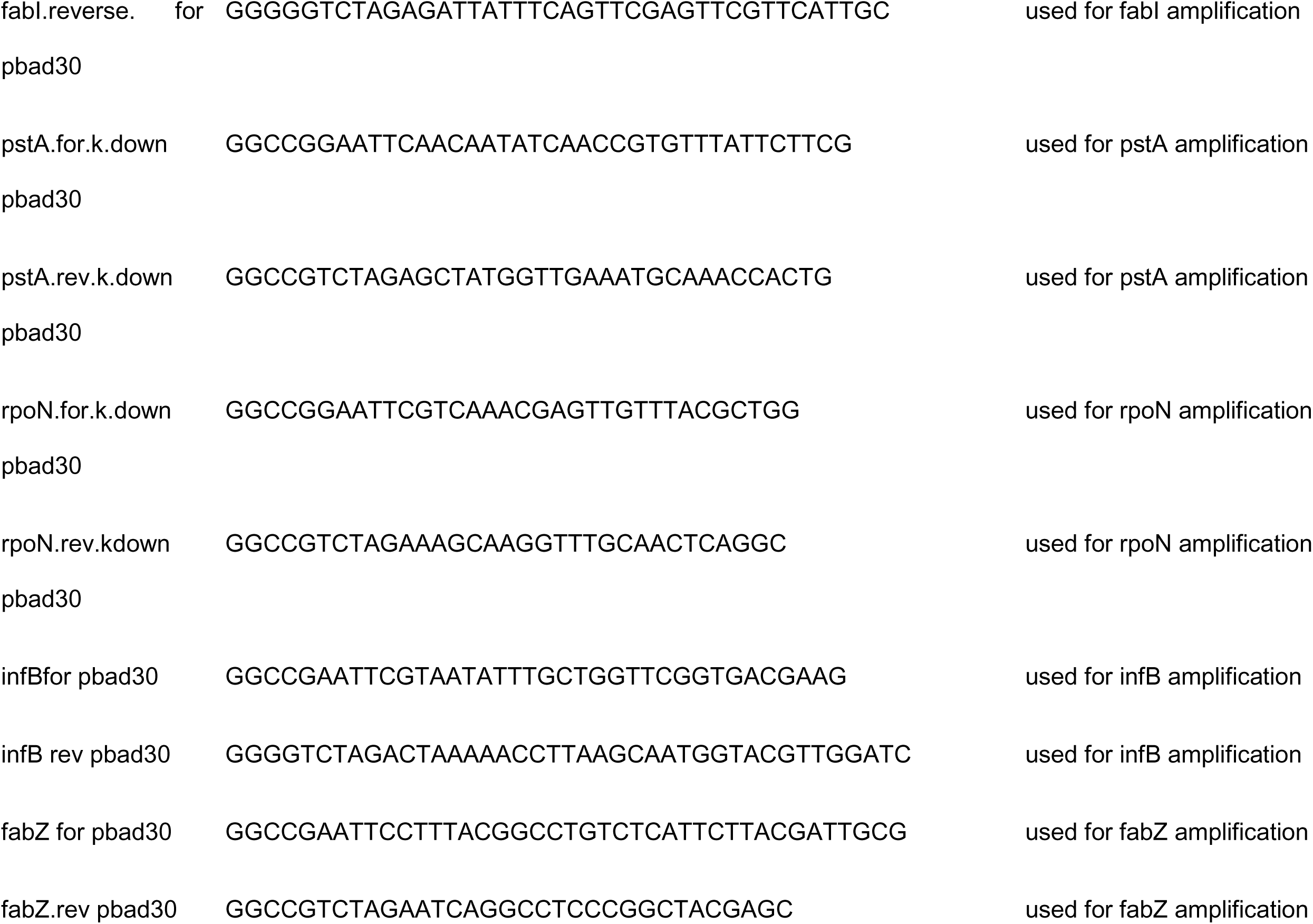
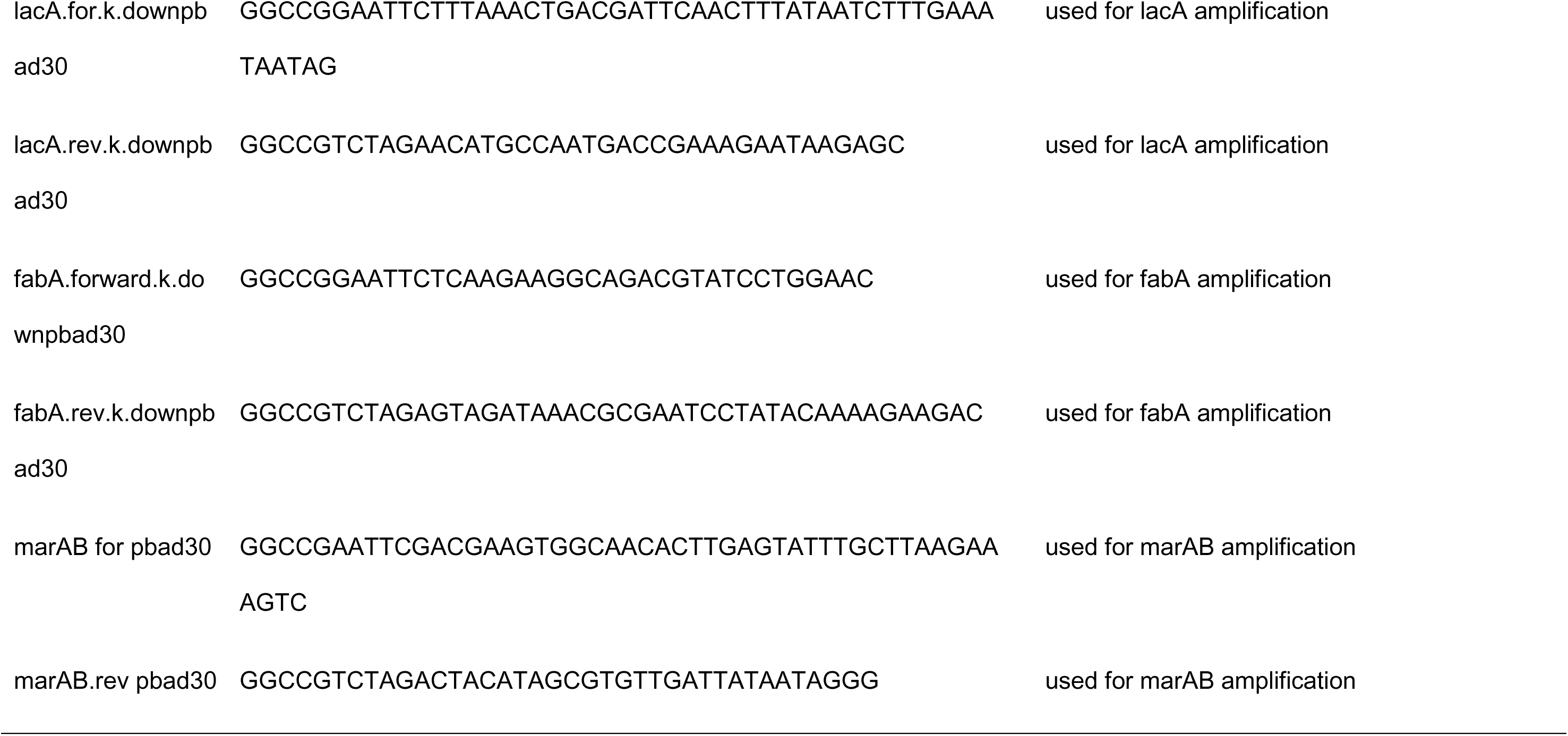
Primer sequences

